# Transient, early, female-specific increase in cortical glial fibrillary acidic protein distribution in the Syrian hamster model of mild peripheral COVID-19

**DOI:** 10.1101/2025.04.08.647811

**Authors:** Mohammadreza Rahmani Manesh, Leigh E. Wicki-Stordeur, Nicole S. York, Robert Vendramelli, Bryce Warner, Haley A. Vecchiarelli, Luke Rainier-Pope, Mohammadparsa Khakpour, Lucas R. Bennouna, Marie-Ève Tremblay, Darwyn Kobasa, Leigh Anne Swayne

## Abstract

**Background:** Mild-moderate respiratory COVID-19 is commonly associated with a range of neurological symptoms. The mechanisms linking this peripheral disease to cognitive symptoms are thought to include heightened circulating cytokines and other inflammatory mediators resulting in a leaky blood-brain barrier and increased neuroinflammation (i.e., inflammation taking place in the brain). This can lead to aberrant synaptic transmission and cognitive dysfunction. A key component of neuroinflammation is the reactivity of astrocytes, in a process termed ‘astrogliosis’, associated with altered morphology, proliferative capacity, gene expression, and function. Accumulating evidence suggests astrogliosis likely occurs in mild-moderate COVID-19; however, there has been limited investigation. In this study, we quantified changes to astrocytes in a Syrian hamster model of mild-moderate respiratory COVID-19.

**Methods:** We used an intranasal inoculation model to produce mild-moderate respiratory COVID-19 in 8–10-week-old male and female Syrian hamsters. We extracted brains at 1-, 3-, 5-, 7-, and 31-days post-inoculation and from uninfected controls, and immunolabelled brain sections with astrocyte- (GFAP and SOX9) and neuron-specific (NEUN) markers. We captured tiled confocal micrographs of entire brain sections and analyzed the resulting signals from five regions of interest: cortex, corpus callosum, hippocampus, third ventricle, and dorsal striatum.

**Results:** To systematically quantify cell-type-specific labelling for astrogliosis markers, we first developed an unbiased pipeline. We found a transient increase in GFAP signal density in female hamster, specifically in the cortex at 3 days post-inoculation. There were no corresponding changes noted in astrocyte (SOX9), neuron (NEUN) or total cell (Hoechst) numbers. Moreover, there were no changes in male hamsters at any timepoint in any region of interest.

**Conclusions:** Our findings provide the first spatiotemporal insight into astrogliosis in a hamster model of mild-moderate respiratory COVID-19. We identified a transient and sex-specific increase in GFAP signal density, indicative of astrogliosis. Our findings contribute to the literature surrounding sex differences in (neuro)immune responses and add to the growing body of COVID-19 literature, in which sex-specific outcomes are apparent in both human patient populations and rodent experimental models.

## Background

Coronavirus disease 2019 (COVID-19) is an infectious disease caused by severe acute respiratory syndrome coronavirus 2 (SARS-CoV-2) infection and the mild to moderate form of the disease is commonly associated with respiratory tract symptoms (reviewed in (1)). Additionally, COVID-19, presenting as asymptomatic to severe in terms of the respiratory disease, can be associated with a range of mild to severe neurological symptoms (reviewed in (2–5)), the development and severity of which tend to be directly correlated with the severity of COVID-19 respiratory disease (reviewed in (3)). Notably, a spectrum of COVID-19 symptoms, including neurological symptoms, can persist or even develop after the acute infection is cleared (commonly referred to as ‘long COVID’) even with asymptomatic to mild COVID-19 disease (reviewed in (2,4,6)). Common cognitive neurological symptoms associated with long COVID include ‘brain fog’ (colloquial term for a variety of symptoms associated with cognitive impairment, such as difficulty concentrating, confusion, and fatigue (7)), anxiety, and depression (reviewed in (2,4)). Importantly, these neurological symptoms affect women at a disproportionate rate (e.g. (8,9)).

The cellular and molecular mechanisms underlying the development of COVID-19- associated cognitive symptoms are likely to involve neuroinflammatory processes ((10,11); reviewed in (4,12,13)). *In vitro*, SARS-CoV-2 is taken up into various types of brain cells; however, *in situ* brain tissue detection of SARS-CoV-2 is inconsistent, and there is no clear evidence of the presence of whole virus or viral replication in the brain (discussed in (14)). There is strong evidence of increased circulating cytokines, which in other inflammatory disorders has been associated with the development of neuroinflammation, in part due to increased blood-brain-barrier permeability (reviewed in (4,15)). Given the strong evidence of blood-brain barrier disruption in COVID-19 (reviewed in (15)), a similar mechanism has been proposed for triggering neuroinflammation in the context of COVID-19, whereby a relatively leaky blood-brain barrier provides a route for the increased circulating peripheral cytokines into the brain, thereby triggering inflammatory signaling mechanisms in glial cells. More recent evidence (14) suggests the SARS-CoV-2 spike protein may accumulate and persist in the skull-meninges-brain axis of human COVID-19 patients, possibly increasing inflammation locally, thereby enhancing the movement of inflammatory molecules into the brain parenchyma. Supporting this ‘peripheral to neuro’ inflammation mechanism for mild COVID-19-associated neurological symptoms, neuroinflammation was observed in a mouse model of mild respiratory COVID-19 (16), in which human ACE2 was delivered via adeno-associated virus to the mouse trachea and lungs to elicit a mild respiratory infection. These mild respiratory COVID-19 mice exhibited microglial reactivity in the cortex, corpus callosum, and hippocampus, as well as impaired hippocampal neurogenesis, decreased oligodendrocyte density, as well as increased cytokine and chemokine levels in the cerebrospinal fluid (16). Notably, many of the observed changes persisted long after the respiratory infection cleared. Similarly, the golden, or Syrian hamster (*Mesocricetus auratus*), which endogenously expresses ACE2 and model mild to moderate respiratory COVID-19, exhibited increased olfactory bulb microglial density (17,18) along with inflammatory cytokine and microglial reactivity marker transcript expression (17). While no overt histological changes were observed in the brain (19), bioinformatic (gene ontology; GO) analysis of transcriptomic changes revealed changes in GO terms associated with neuro-immune responses, including a female-specific enrichment of the general GO term “inflammatory response” (19). Together these findings support neuroinflammation in the context of mild to moderate COVID-19, with potential sex differences, and directly implicate microglia. In addition to changes in microglia, astrogliosis was also proposed to underlie COVID-19-associated cognitive dysfunction (e.g., as reviewed in (20)) given the important role that astrocytes play in synapse stability (e.g., as reviewed in (21,22)). Astrogliosis has primarily been investigated in the context of critical acute COVID-19 in post-mortem neuropathological studies (reviewed in (23)).

Whether astrogliosis occurs in mild to moderate COVID-19 at the acute or chronic stage (i.e., following resolution of the acute infection) is relatively understudied. This is an important knowledge gap given that neurological symptoms and neuroinflammation occur in mild COVID-19 at the acute stage as well as after the virus has cleared (e.g. (16)).

Comprehensive characterization of changes in the brain with mild respiratory COVID-19 will enable improved mechanistic understanding and management of neurological symptoms. Astrogliosis was recently investigated in human patients with ongoing depressive and cognitive symptoms following mild to moderate COVID-19 using a non-invasive positron-emission tomography strategy (24). Importantly, this groundbreaking work supports the presence of astrogliosis in the context of mild COVID-19 but provides limited spatiotemporal insight.

To quantify astrogliosis with spatiotemporal resolution, and explore potential sex differences, we used the Syrian hamster model. Intranasal inoculation of Syrian hamsters produces mild to moderate respiratory COVID-19 that shares many similarities with human disease in terms of host responses and time to resolution ((17,25,26)(reviewed in (27)). Measurement of astrogliosis is challenging because it can involve a plethora of changes, including marked alterations in both the density and morphology of astrocytes (reviewed in (28–30)). Risk mitigation protocols, such as long-term fixation, surrounding manipulation of COVID-19 tissues further compound these challenges. Quantification of astrogliosis in the Syrian hamster model of COVID-19 required the development and validation of a systematic imaging pipeline. We used our custom pipeline to measure the distribution of the astrocyte-enriched glial fibrillary acidic protein (GFAP) ((31,32); reviewed in (33)), and density of astrocyte-enriched SRY-box transcription factor 9 (SOX9)-positive cells (34) from tiled confocal images of full coronal brain sections from male and female hamsters, at 1-, 3-, 5-, 7-, and 31- days post-inoculation (dpi), and in uninfected controls. We discovered a significant transient increase in GFAP distribution at 3-dpi in the cortex of female animals. No changes were observed in SOX9, or other markers outlined below. Together our results reveal mild/partial, transient astrogliosis during the acute phase of infection, in female animals, that resolves in the long term.

## Methods

### Virus preparation

The SARS-CoV-2 (Canada/ON-VIDO-01/2020; EPI_ISL_425177) virus used was previously isolated from a positive patient sample in 2020 and stocks of the virus were grown in VeroE6 cells. All virus used for *in vivo* experiments was from passage 2 as described previously and were titered by TCID50 assay (35).

### Animals

All animal work was conducted in compliance with the guidelines established by the Canadian Council on Animal Care, as approved by the Animal Care Committee at the Canadian Science Center for Human and Animal Health (CSCHAH: animal use document H-20-006). 8-10-week-old Syrian hamsters (*Mesocricetus auratus*; Charles River Laboratories, Wilmington, Massachusetts, USA) were used in this study. Hamsters were monitored by registered Animal Health Technicians throughout the experiment and were provided food and water *ad libitum*. Animals were group housed under controlled laboratory conditions, including a 12-hour light/dark cycle, a temperature range of 21–22°C, and humidity levels of 30–40%. Animals were acclimatized for 7 days before the beginning of all experiments.

For SARS-CoV-2 infection, hamsters were inoculated intranasally with 10⁵ TCID50 of the original VIDO (Wuhan-like)-strain under isoflurane anesthesia in the Containment Level 4 laboratory at the National Microbiology Laboratory of the CSCHAH. A total of 100 μL of the viral inoculum was divided equally between nostrils. Uninfected controls received no inoculum. Animals were monitored and weighed daily, and those that reached euthanasia criteria or experimental endpoints (1-, 3-, 5-, 7-, and 31-dpi) were anesthetized with isoflurane and euthanized by exsanguination and cervical dislocation. Brains were removed and fixed for a minimum of 30 days in 10% formalin before processing according to Standard Operating Protocol for removal of tissues from the high containment laboratory.

### Determination of viral burden in tissues

Nasal turbinates and distal lungs were collected from all animals and flash frozen at -80°C until further analysis for live virus titers. Tissues were homogenized in 1 mL viral growth media (MEM supplemented with 1% Bovine Growth Serum (BGS, Cytivia) and penicillin and streptomycin (100 Units/mL and 100 µg/mL respectively; Gibco)), along with 5 mm sterile stainless steel beads using a Bead Ruptor Elite Tissue Homogenizer (Omni). Homogenates were clarified by centrifugation at 1500 x g for 10 minutes, and ten-fold dilutions of tissue homogenates were made in viral growth media. Dilutions were added to Vero-TMPRSS2 cells (BPS Bioscience, Cat# 78081) cells in triplicate wells, and cytopathic effect was read 96 hours post-infection. TCID50 values per gram of tissue were calculated using the Reed and Muench method (36).

### Immunohistochemistry

All procedures were performed by personnel blinded to the experimental conditions. Highly fixed brains were sectioned on a vibratome (Leica VT1000, Leica Camera, Wetzlar, Germany) into 50 μm coronal slices in Dulbecco’s PBS (DPBS; catalogue #14190250, Thermo Fisher Scientific, Waltham, Massachusetts, USA). Sections were stored at -20°C in cryoprotectant (DPBS, 30% ethylene glycol, 30% glycerol) until immunolabelling occurred. Sections were chosen for immunolabelling based on anatomical landmarks identified using the Allen Mouse Brain Atlas (https://mouse.brain-map.org/experiment/thumbnails/100142143) and the Golden Syrian hamster atlas (37); those chosen for further analyses included slices containing the dorsal hippocampus (equivalent to mouse bregma -1.255 mm to -2.355 mm) and the lateral ventricles (equivalent to mouse bregma 1.145 mm to 0.020 mm). All immunolabelling experiments included equal sections from each experimental group (control and 1-, 3-, 5-, 7-, and 31- dpi). Chosen sections were washed three times for 5 minutes each in DPBS, permeabilized with 1% Triton-X in DPBS for 30 minutes at room temperature, and blocked with 10% normal donkey serum (catalogue #017-000-121, Jackson ImmunoResearch Labs, Pennsylvania, USA) in 0.5% Triton-X in DPBS for 1 hour at room temperature. Primary antibody incubation was performed at 4°C with shaking for two nights, with antibodies diluted in 5% normal donkey serum and 0.5% Triton-X in DPBS. Primary antibodies used were anti-SOX9 (catalogue #82630S, 1:200; Cell Signaling Technologies, Danvers, Massachusetts, USA), anti-GFAP (catalogue #13-0300, 1:200; Invitrogen, Waltham, Massachusetts, USA), and anti-NeuN (catalogue #MAB377, 1:500; Millipore Sigma, Burlington, Massachusetts, USA). Sections were washed three times for 5 minutes in DPBS then incubated with secondary antibodies in 5% normal donkey serum and 0.1% Triton-X in DPBS for 1 hour at room temperature. Secondary antibodies used were donkey anti-Rat IgG (H+L) Alexa Fluor 488 (catalogue #712-545-150; 1:300; Jackson ImmunoResearch Laboratories), donkey anti-Mouse IgG Alexa Fluor 568 (catalogue #A-10037; 1:300; Thermo Fisher Scientific), and donkey anti-Rabbit IgG Alexa Fluor 647 (catalogue #A-31573; 1:300; Thermo Fisher Scientific). Secondary antibodies were centrifuged at 4°C for 5 minutes prior to use. Hoechst 33342 (catalogue #62249, 1:1000; Thermo Fisher Scientific) was used as a nuclear counterstain. Sections were washed four times for 5 minutes each in DPBS then mounted onto glass slides with ProLong Gold antifade reagent (catalogue #P36934, Thermo Fisher Scientific). Slides were cured at room temperature protected from light for at least 24 hours, then stored at -20°C.

### Confocal immunofluorescence imaging

Samples were imaged using a Leica TCS SP8 line scanning confocal microscope with Leica Application Suite Software version 3.1.3.16308. Using the tiling feature, entire brain sections were imaged with a 20X (NA 0.7) air objective at 512 x 512-pixel resolution. The resulting images were processed and analyzed using a custom pipeline, the development of which is described in the Results section.

### Scanning electron microscopy

Fifty μm coronal sections containing the primary somatosensory barrel cortex and dorsal hippocampus (equivalent of mouse Bregma -0.95 mm to -1.91 mm) were selected and additionally post-fixed in 0.6 % glutaraldehyde for 2 hours to preserve ultrastructure, followed by three, ten-minutes washes in PBS, prior to long-term storage in cryoprotectant.

Slices were additionally post-fixed and embedded for ultrathin sectioning and scanning electron microscopy (SEM) as previously described (38). Briefly, following five, five-minute washes with PBS to remove cryoprotectant, at room temperature, slices were additionally post-fixed in 2 % osmium tetroxide (Electron Microscopy Sciences, catalogue #19191) containing 3 % potassium ferrocyanide (Millipore Sigma, catalogue #P9387) in phosphate buffer (PB) for one hour. Slices were then washed subsequently with PB, PB:double distilled water (ddH_2_O) mix and ddH_2_O, each for five minutes, followed by a twenty-minute incubation in filtered 1 % thiocarbohydrazide (Millipore Sigma, catalogue #223220) in ddH_2_O. Slices were then washed three times five minutes in ddH_2_O followed by a thirty-minute incubation in 2 % osmium tetroxide in ddH_2_O before five, five-minute washes in ddH_2_O and dehydrated in increasing concentrations of ethanol (five-minute washes, each: 35 %, 35 %, 50 %, 70 %, 80 %, 90 %, 100 %, 100 %, 100 %). Slices were treated with propylene oxide (Millipore Sigma, catalogue #110205) in three times five-minute washes before they were impregnated in Durcupan resin (Millipore Sigma, catalogue #44610) overnight. The next day, sections were mounted between ACLAR embedding films (Electron Microscopy Sciences, catalogue #44610) and polymerized at 55 °C for 72-96 hours.

The barrel cortex was excised from the embedding films and re-embedded on a resin block, sectioned at 73 nm thickness using a Leica ARTOS ultramicrotome. Ultrathin sections were imaged on a Zeiss Crossbeam 350 SEM running ATLAS software. The images were acquired using the ESB and SE2 detectors of the microscope using a 5 mm working distance, at a voltage of 1.4 kV and current of 1.2 nA. Images were acquired at 25 and 5 nm/pixel, tiled together and exported as tiff files.

### Experimental design and statistical analyses

Equal numbers of male and female hamsters were collected for each experimental group (N = 5 per sex). Depending on the specific brain region, N = 3-5 biological replicates met minimum tissue quality levels, which was somewhat variable as no sharp objects could be used in brain extraction as per approved protocols at the high containment laboratory.

Researchers were blinded to the identity of the experimental groups from tissue sectioning until after quantification of immunolabelled confocal micrographs. All data are presented as mean ±SEM. Hamster weight data were analyzed by mixed-effect analysis (due to decreasing sample size (N) over time) with post-hoc Šídák’s multiple comparisons test.

Pipeline outputs from immunolabelled confocal micrographs of male and female hamsters were analyzed separately by one-way ANOVA with post-hoc Dunnett’s comparison to control uninfected of the same sex. Data were analyzed using GraphPad Prism version 10.4.1 (GraphPad Software) and RStudio v4.4.0 (RStudio). Statistical significance was determined by p < 0.05 in all tests. Significance is denoted as *p < 0.05, **p < 0.01, ***p < 0.001, ****p < 0.0001. Statistical results can be found in corresponding Figure Legends as well as the Supplementary Materials.

## Results

### Syrian hamster models mild COVID-19 following intranasal SARS-CoV-2 inoculations

To model mild COVID-19, we inoculated Syrian hamsters with the ancestral Wuhan-like strain of SARS-CoV-2 and sacrificed the animals at 1-, 3-, 5-, 7-, and 31-dpi (Figure 1A). We quantified infectious virus in the nasal turbinates and distal lungs to monitor infection status, and recorded animal weight to track sickness and recovery behaviours. Viral particle numbers peaked early in nasal turbinates and distal lung at 1-dpi in both male and female animals and gradually declined to virtually undetectable levels by 7-dpi (Figure 1B), indicating resolution of the active infection period. No differences in viral load were noted between males and females. Accordingly, all animals exhibited weight loss between 1-7-dpi then re-gain weight up to 20-dpi (Figure 1C). As previously reported (e.g., (19,39)), female animals showed more rapid and greater weight re-gain than males (mixed-effect analysis: Time (p < 0.0001), Sex (p < 0.0001), Time x Sex (p < 0.0001); see Table S1 for post-hoc Šídák’s multiple comparisons between males and females). Together these data suggest that while the acute infection period did not differ between sexes, female animals recovered more quickly than males.

**Figure 1.**
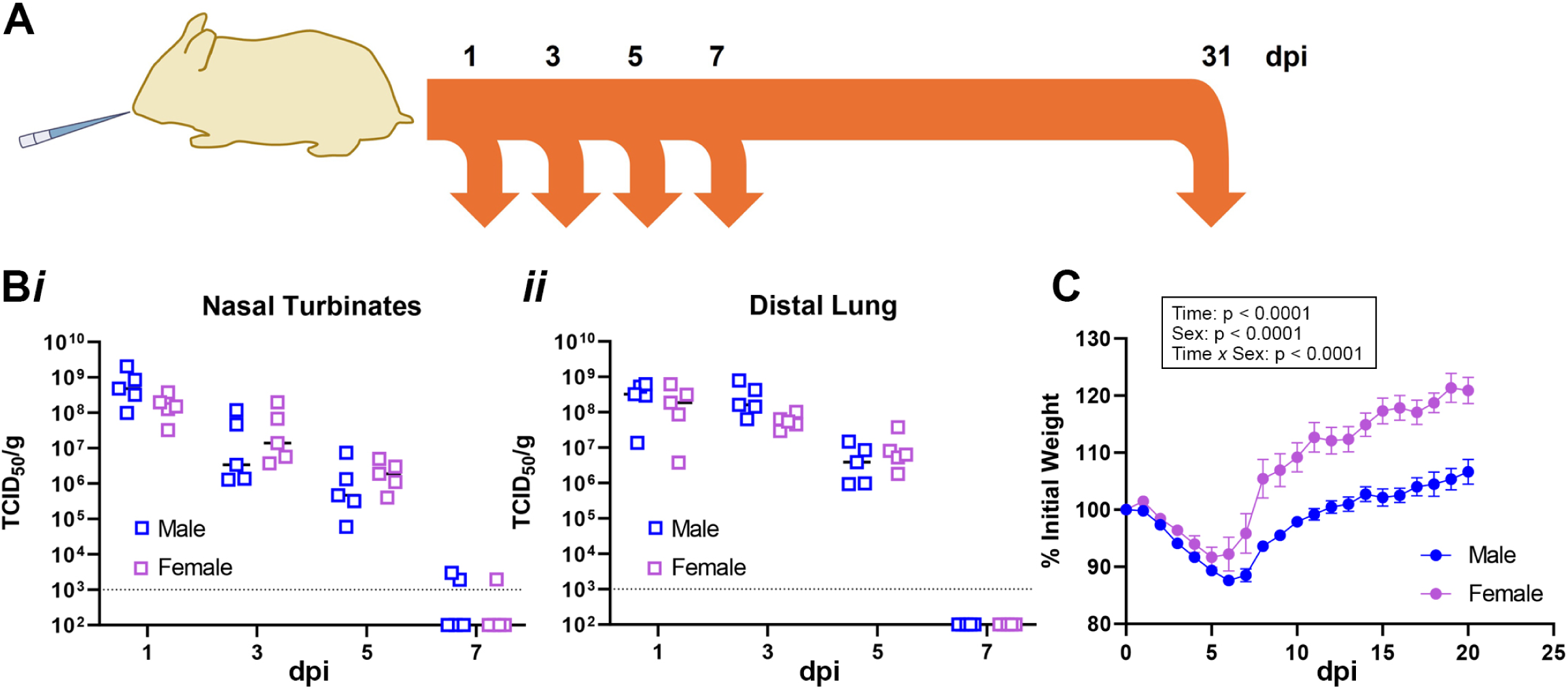
Syrian hamsters model mild-moderate COVID-19 following inoculation with SARS-CoV-2 virus. (A) Experimental model. Equal numbers of male and female 8–10-week-old Syrian hamsters were inoculated intranasally with SARS-CoV-2. Animals were collected at 1-, 3-, 5-, 7-, and 31-days post-inoculation (dpi), along with untreated controls. Equal numbers of male and female animals were collected at each time point (N=5 each). (B) Viral particles were quantified from nasal turbinates (*i*) and distal lung (*ii*) at 1-, 3-, 5-, and 7-dpi. The viral load peaked early and was virtually undetectable by 7-dpi in both regions. No differences were noted between sexes. (C) Inoculated animals exhibited rapid weight loss between 1-7-dpi, then re-gained weight. Female animals had a more rapid and substantial weight re-gain than males (see Table S1).

### Female hamsters show an acute and transient astrocytic response following peripheral SARS-CoV-2 infection

To investigate the brain cellular response to mild peripheral COVID-19 infection, we labelled full brain sections from control and 1-, 3-, 5-, 7-, and 31- dpi Syrian hamsters with cell type specific antibodies (astrocytes: GFAP, SOX9; neurons: NEUN) and a total nuclear marker (Hoechst 33342). We captured tiled confocal images of the full brain sections and analyzed five regions of interest (ROIs; Figure 2), including cortex, corpus callosum, hippocampus, third ventricle, and dorsal striatum. We further developed a custom MATLAB analysis pipeline to quantify signals from this confocal dataset (Figures 3 and S1). The MATLAB codes are freely available on GitHub: https://github.com/SwayneLab/Quantitative-analysis-of-astrocyte-properties-in-a-Syrian-hamster-model-of-COVID-19.

**Figure 2.**
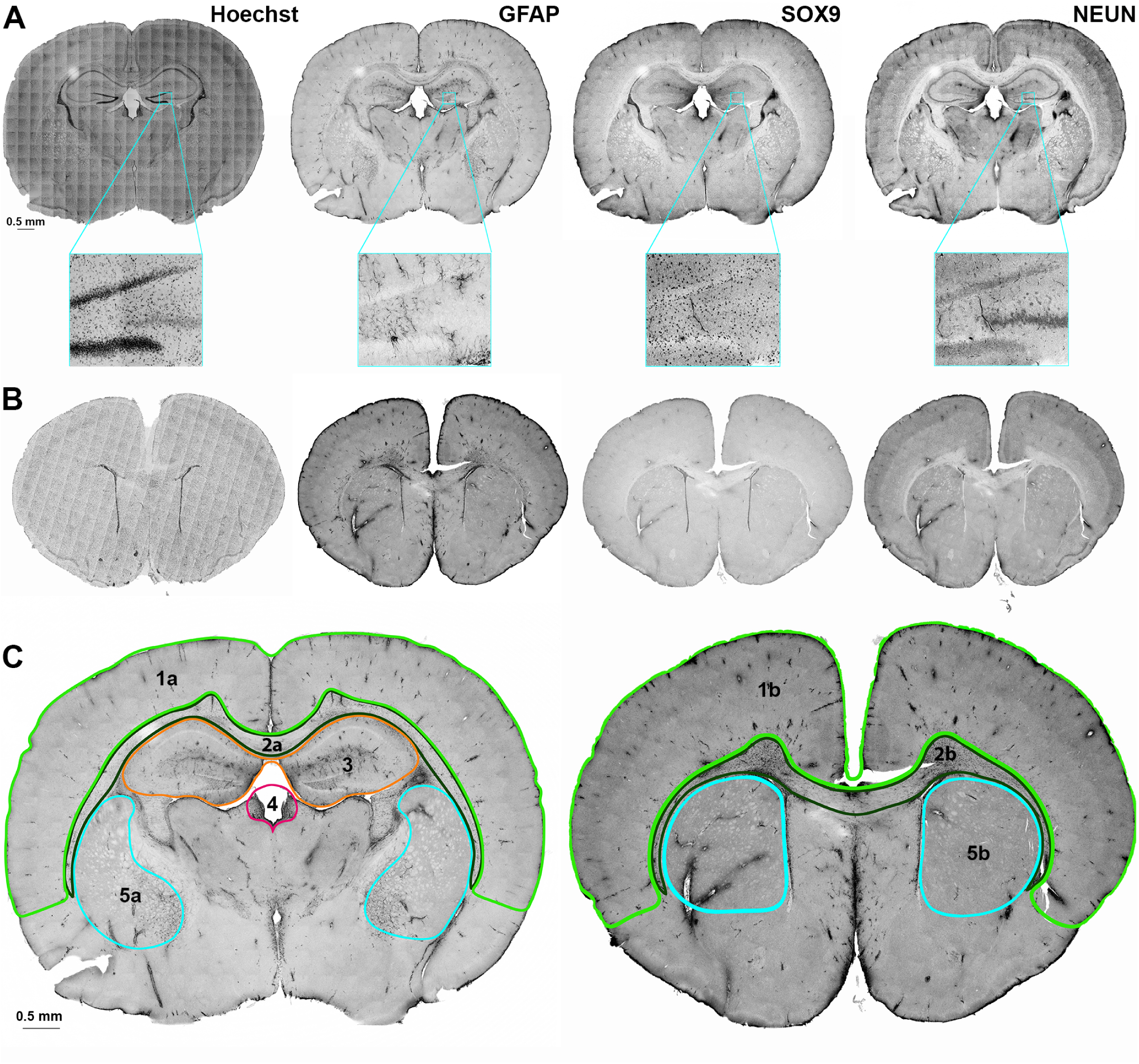
Immunolabelling of full hamster brain sections with cell-type specific markers. Coronal brain slices from infected and control hamsters were labelled with astrocyte-specific GFAP and SOX9 antibodies, neuron-specific NEUN antibody, and the general nuclear stain Hoechst 33342. Slices were selected based on anatomical landmarks, specifically (A) those containing dorsal hippocampus and (B) those containing lateral ventricles. Representative tiled confocal micrographs of (left to right) Hoechst, GFAP, SOX9 and NEUN are depicted, with insets showing zoomed in regions. Scale bars are 0.5 mm. (C) Tiled confocal micrographs of GFAP signal from example coronal brain slices with our regions of interest (ROIs) outlined and numbered as follows: 1a,b, cortex; 2a,b, corpus callosum; 3, hippocampus; 4, third ventricle; 5a,b, dorsal striatum. Scalebar is 0.5 mm. This figure was previously included in the MSc thesis of MRM ((68); link: https://dspace.library.uvic.ca/items/d7256e07-eee8-4b45-80e7-3c4e4404c299)

**Figure 3.**
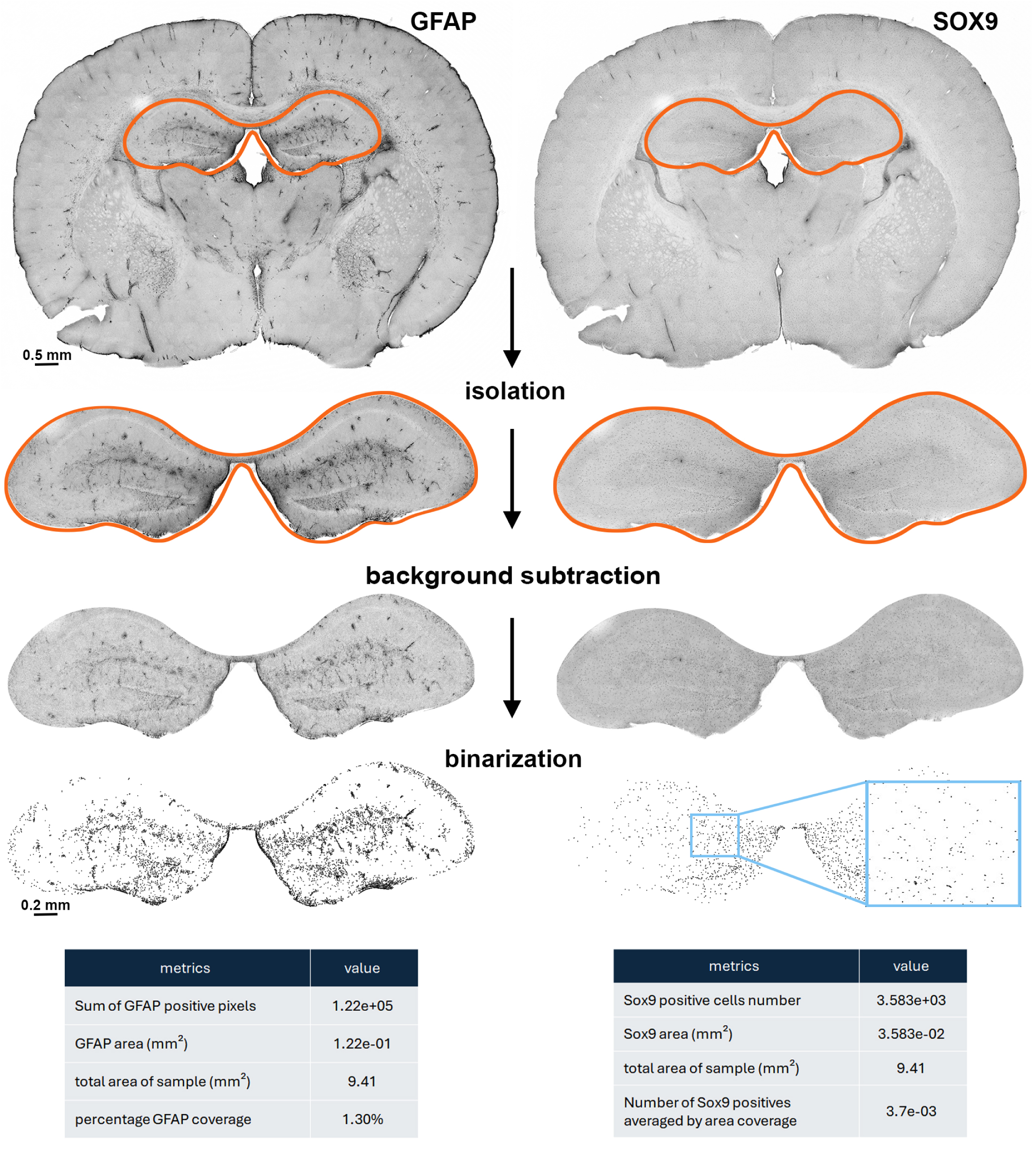
Quantitative analysis pipeline workflow in hamster brain sections immunolabelled for astrocyte markers GFAP and SOX9. (Top) Representative tiled and stitched raw confocal immunofluorescence micrographs of GFAP (left) and SOX9 (right) signals. We first isolate/define the precise region of interest from the full brain slice and determine its area. Next, we improve the signal to noise ratio through background subtraction. Finally, the GFAP and SOX9 signals are converted to a black on white binary format. The pipeline outputs for GFAP and SOX9 (bottom) are normalized to the ROI area, giving signal density for GFAP, and positive cell density for SOX9 as the final read outs. This figure was previously included in the MSc thesis of MRM ((68); link: https://dspace.library.uvic.ca/items/d7256e07-eee8-4b45-80e7-3c4e4404c299)

We first manually isolated our ROIs from each tiled image. Note that the ventricular zone was carefully excluded from corpus callosum and striatum ROIs to avoid potentially confounding ventricular zone neural precursor cell signals. We improved signal clarity and reduced noise by applying background subtraction using a morphological opening function (40,41), and a Gaussian filter (41,42). We then used entropy thresholding segmentation to isolate positive signals through binary conversion (43,44). For a full account of our comparisons of various thresholding techniques, please refer to the Supplemental Materials, Tables S2, S3 and Figure S1. Finally, we calculated the ROI area using MATLAB’s "numel" function to determine density values (41). GFAP and Hoechst signal outputs were pixel density, while SOX9 and NEUN outputs were positive cell density. Based on manual counts of 100 positive cells from different brain regions, we excluded objects smaller than 5 pixels (5 μm) for SOX9 and 7 pixels (7 μm) for NEUN cell density counts from our pipeline. The pipeline steps and outputs for GFAP and SOX9 are depicted in Figure 3, while those for Hoechst and NEUN are shown in Figure S2.

To probe for evidence of astrogliosis, we examined GFAP and SOX9 signals (Figure 4A, B), which give an indication of astrocyte reactivity/process density and astrocyte nuclei numbers, respectively ((31,32,34); reviewed in (33)). Using our custom analysis pipeline, we found an increase in GFAP signal density in the cortex of female hamsters at 3-dpi compared to controls (Figure 4C and see Table S4 for statistics). There was a similar, non-significant increase in GFAP density in the hippocampus (Figure 4D) and corpus callosum (Figure S3A, Table S4) of 3-dpi females compared to controls. There were no changes in GFAP density found in any ROI within males (Figures 4C,D and S3, Table S4). There were no differences in raw GFAP distribution values between control male and female hamsters (data not shown). Notably, there were no changes in SOX9-postive cell density in male or female hamsters (Figure S4, Table S5). Ultrastructural examination of representative control and 3-dpi female cortices using scanning electron microscopy showed lysosomal accumulation and potentially altered mitochondrial cristae structure in astrocytes occupying satellite positions onto neurons at 3-dpi (Figure 5).

**Figure 4.**
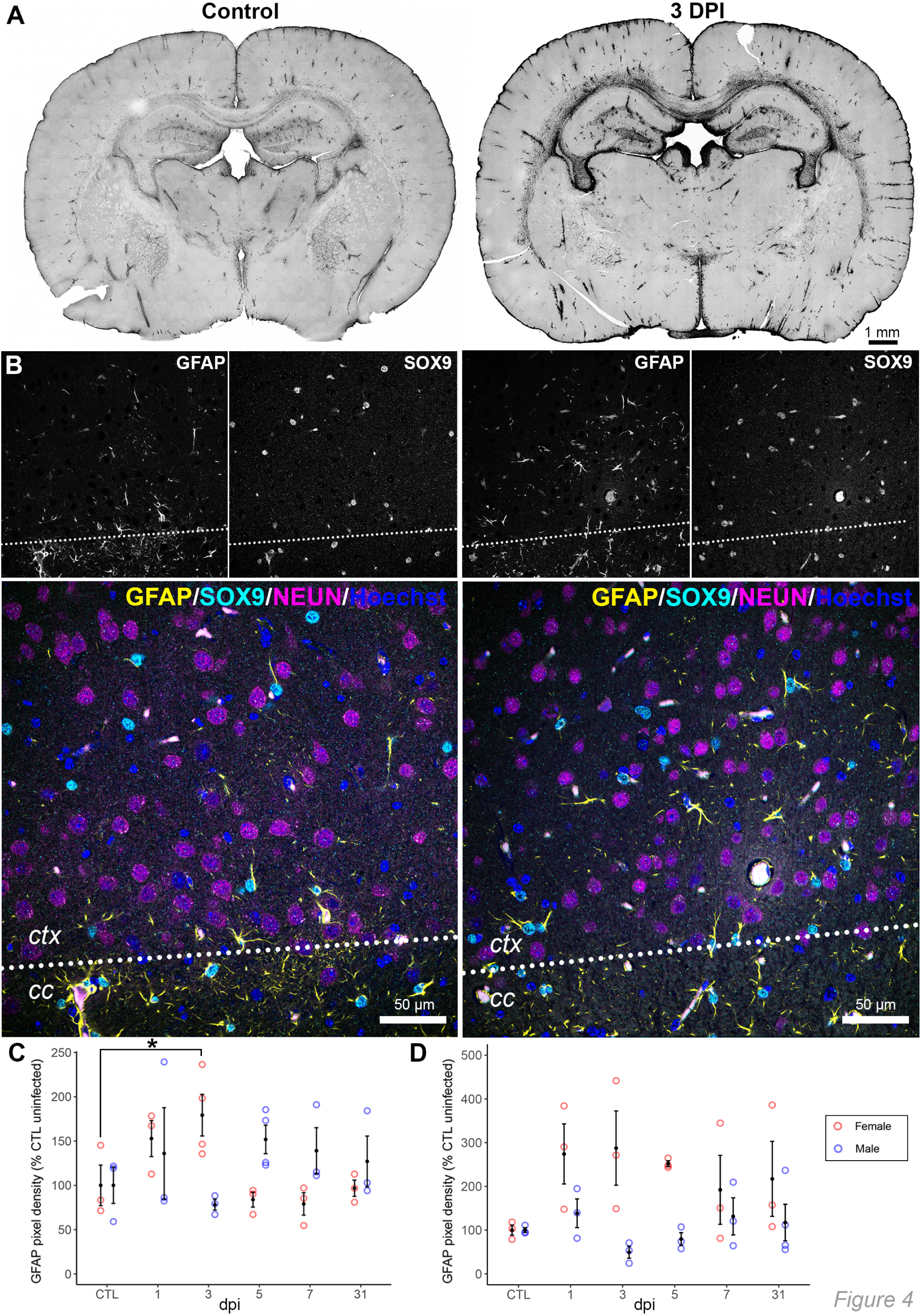
Signal density of the astrocyte marker, GFAP, increases in female hamster brains shortly following peripheral SARS-CoV-2 infection. (A) Representative tiled confocal micrographs showing GFAP signal from full brain slices of uninfected control (left) and 3-dpi (right) female hamsters. Scale bar is 1 mm. (B) Representative high magnification confocal micrographs from uninfected control (left) and 3-dpi (right) female hamsters. Images are from deep cortical layers (*ctx*) at the border of the corpus callosum (*cc*, dotted line). Individual GFAP and SOX9 signals are shown in greyscale, with merged GFAP/SOX9/NEUN/Hoechst micrographs shown below. Scalebars are 50 µm. Quantification of GFAP signal density in the cortex (C) and hippocampus (D) of male and female hamsters. There is a significant increase in GFAP signal density within the cortex specifically in female hamsters at 3-dpi (*) compared to uninfected controls (CTL). A similar, yet non-significant, increase is noted in the hippocampus of 3-dpi females. No changes were noted in males. For quantification of GFAP signal density within other ROIs, see Figure S3, and Table S4 for statistical outcomes.

**Figure 5.**
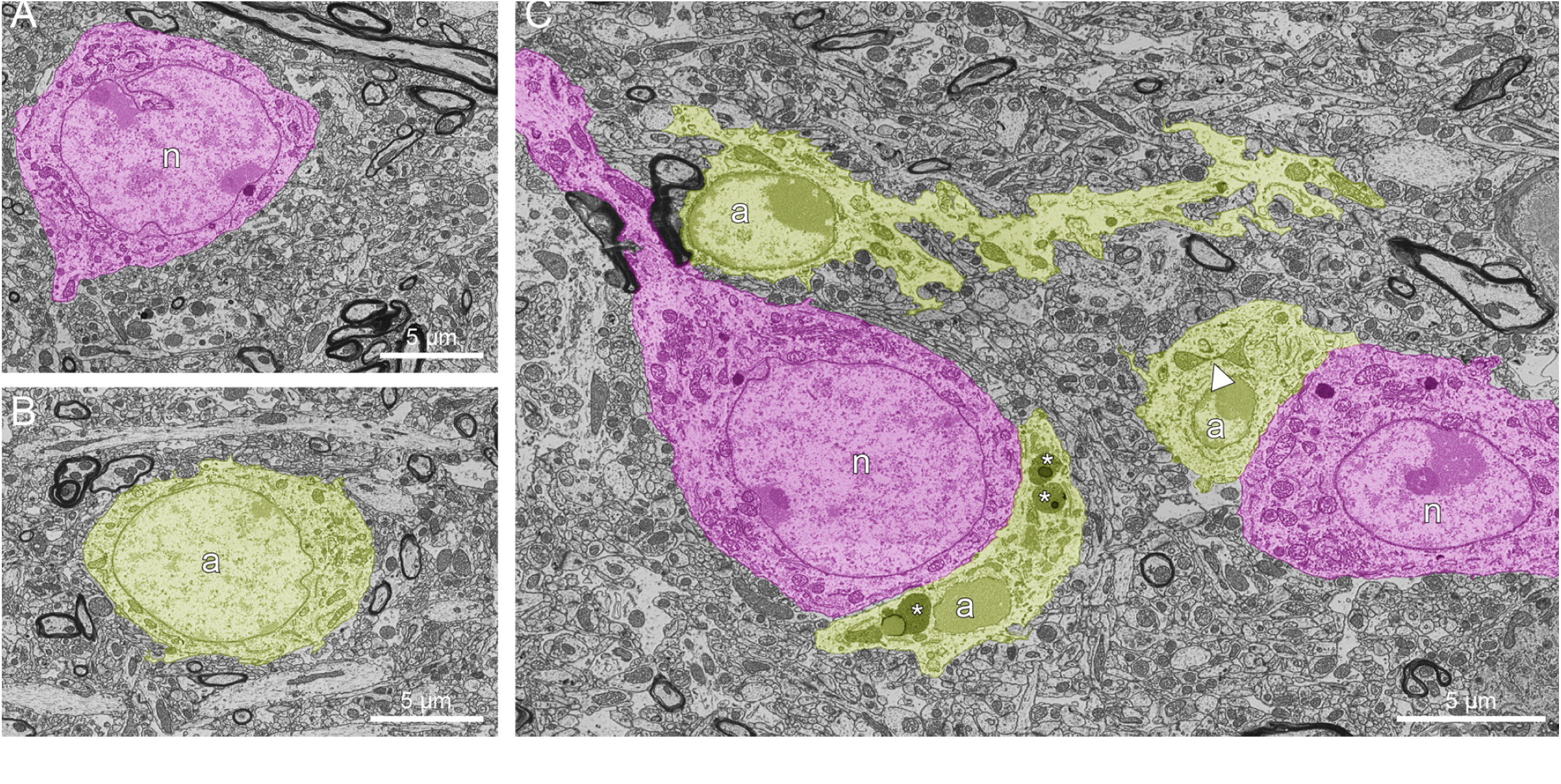
Scanning electron microscopy examples of astrocyte ultrastructure in the 3-dpi female cortex. Example scanning electron micrographs of (A) neurons (n) and (B) astrocytes (a) from control uninfected female cortex. (C) Example scanning electron micrograph from 3-dpi female cortex. The astrocytes show lysosomal accumulation (*) and potentially altered mitochondria (arrowhead) in astrocytes occupying satellite positions to neurons. Scale bars 5 µm.

We further probed for changes in neuron or total cell density within our samples using the neuron-specific nuclear marker, NEUN, and the general nuclear stain, Hoechst 33342. There were no changes in either NEUN-positive cell density (Figure S5, Table S6) or Hoechst pixel density (Figure S6, Table S7) in SARS-CoV-2-infected samples compared to controls.

Overall, our data suggest there is an early and transient astrocytic response to mild peripheral SARS-CoV-2 infection. This occurs without changes to astrocyte, neuron, or total cell numbers.

## Discussion

Here, we developed a robust pipeline for quantification of astrocyte density, based on SOX9 immunoreactivity and astrocytic GFAP distribution, to investigate brain-wide astrogliosis in mild respiratory COVID-19 with improved spatiotemporal resolution in both sexes. Our primary significant observation was a transient increase in GFAP distribution in the cortex at 3-dpi, in females only. Similarly, we observed an increase in GFAP distribution in the hippocampus and corpus callosum at 3-dpi in females only, albeit these were non-significant. Despite these differences in GFAP distribution, no changes in astrocyte density were observed. This is perhaps not surprising, given that astrocyte proliferation is limited in inflammatory contexts even when there are increases in GFAP (reviewed in (30)). Together, these data suggest mild, respiratory COVID-19 elicits early astrogliosis in the cortex of females and indicate the female hippocampus and corpus callosum may warrant further investigation.

In terms of the affected regions, both cortex and hippocampus are highly vascularized (albeit the cortex much more so than the hippocampus as reviewed in (45)), making the astrocytes therein readily accessible to inflammatory mediator infiltration and the ensuing signalling cascades that can culminate in gliosis (reviewed in (12)). There is evidence that the blood-brain barrier is disrupted in COVID-19 (15), and although this has not yet been established in the context of mild respiratory COVID-19, our results and those of others (e.g., (16–18)) suggest this occurs. Interestingly, SARS-CoV-2 variants may differentially affect blood-brain barrier components and integrity (46,47) and, therefore the downstream astrogliosis. This is a significant consideration given our study was limited to the use of an ancestral Wuhan-like strain of the virus.

In terms of inflammatory mediator involvement, interleukin (IL)1-β, IL-6, and tumor necrosis factor-α (TNF-α), increase substantially in nasal turbinates and/or lung within 2- dpi (and possibly earlier) in the hamster model of COVID-19 (48). These cytokines all elicit astrogliosis (reviewed in (49)). Notably, female-specific brain transcriptional changes associated with the GO term “inflammatory responses” in the Syrian hamster model were observed at 2-dpi (19). Therefore, the timing of the changes we see in GFAP density in females is in line with previous findings in peripheral cytokine increases, as well as female-specific inflammatory changes in the brain. There are considerable sex-differences in astrocyte gene expression (50), and therefore there are several possible mechanisms at play. Notably, astrocytic cytokine release, another measure of astrocyte reactivity known to be triggered by cytokine exposure, was greater in males than females in response to lipopolysaccharide ((51); reviewed in (52)).

Our findings from the female corpus callosum are also notable, given a previous study using a mouse model of mild to moderate COVID-19 found evidence of glial cell activation in the form of microgliosis in this region (16). While the corpus callosum possesses a much lower degree of vascular density than both cortex and hippocampus (53), heightened microglial reactivity can contribute to astrogliosis by increasing local production and release of cytokines and other inflammatory molecules (reviewed in (54,55)). Similarly, increased microglial density and transcriptional markers of microglial reactivity were found in the olfactory bulb of golden (Syrian) hamsters (17,18). The olfactory system is particularly susceptible in both human COVID-19 cases and rodent models of the disease (e.g., (56–59)). It is important to note that we were unable to reliably extract the olfactory bulbs due to limitations on the use of sharp instruments imposed by standard safety procedures in the high biosafety containment lab, and as such this presents a conspicuous ROI for future studies.

Our SEM analysis indicated lysosomal accumulation and potentially altered mitochondria in astrocytes of the 3-dpi female cortex. Lysosomal changes can impair certain aspects of astrocyte function and contribute to astrocyte-mediated neuroinflammation in the context of brain injury or disease states (reviewed in (60)). Intriguingly, GFAP may influence lysosomal function (61,62), and mutations causing GFAP accumulation in astrocytes (Alexander’s disease) result in both lysosomal and mitochondrial defects (63,64). It therefore remains to be seen whether these observations are due specifically to increases in GFAP at 3-dpi or, more generally, to increased inflammation and resulting astrogliosis. Moreover, whether similar alterations occur in male animals, and whether these ultrastructural abnormalities persist or resolve alongside GFAP density metrics in the female cortex are unknown.

Our detection of astrogliosis in the context of mild to moderate respiratory COVID- 19 was not unexpected, given forms of gliosis have also been observed in the context of mild to moderate peripheral COVID-19, in both animal models (e.g., (16–18)) and human studies (e.g., (24,65)), including specifically astrogliosis (24). Consistent with our findings, in the human study, mild to moderate COVID-19 was associated with increased GFAP distribution in all cortical areas assessed, as well as in the hippocampus (24). Interestingly, in contrast to the human study, we did not observe increased GFAP distribution in the striatum, and we did not see long-term changes in astrocytes. These differences in outcomes may in part be attributed to differences in the range of illness severity (relatively narrow in hamsters *vs*. broader in humans, as well as the primary inclusion criteria in the human study being the onset of a new major depressive episode), to the variant of SARS-CoV-2 (ancestral variant in ours *vs.* presence of omicron in the human study) and also to the sensitivity and longevity of the marker (GFAP distribution in our study *vs.* total distribution volume of [11C]SL25.1188, an index of monoamine oxidase B density, in the human study). No effect of sex was observed in the human study, but it is difficult to know whether the study was sufficiently powered to detect such effects given the considerable variability in a range of key parameters.

Our findings support observations made in the earlier human study (24) suggesting that astrogliosis in mild to moderate COVID-19 could be protective. In the human work, the extent of astrogliosis was inversely proportional to symptom severity (24). As hamsters become sick with COVID-19, they rapidly lose weight within 1- to 2-dpi (48), and weight regain therefore acts as a correlate of recovery (48). In our study, female hamsters recovered more quickly, at least using weight regain as a wellness metric (Figure 1C), consistent with previous work by others and us (e.g. (19,39)). Given we saw early astrogliosis in females, and females recovered from illness more quickly, it is reasonable to speculate that our findings support the idea brought forth in the human study, that astrogliosis can be protective. Although it is possible that differences in the immune or neuroimmune/inflammatory response in males and females could play a role in the sex-specific recovery rate (e.g. reviewed in (66), comparable viral titers were observed in nasal turbinate and the distal lung (Figure 1B), therefore males and females appear to clear the virus at a similar rate. Our findings raise the intriguing possibility that the astrocyte changes unique to female hamsters, likely elicited in response to the increase in circulating cytokines as outlined above, could contribute, at least in part, to their improved recovery.

There is precedent for sex differences in astrocyte responses to chronic stress and inflammation (reviewed in (52)), such that astrocytes in males tend to respond by secreting higher levels of inflammatory mediators (51), as noted above, while those in females tend to exhibit greater resilience and adopt a more ramified morphology (67). Although GFAP does not extend through to the fine astrocyte processes, it is present in the astrocyte cell body and proximal processes and, as such, can be viewed as a correlate, albeit an imperfect one, of astrocyte morphology (reviewed in (33)), with the increase in GFAP distribution potentially indicative of a more ramified morphology). Astrocytes are well known to play a crucial role in regulating synaptic plasticity (e.g., as reviewed in (21,22)); therefore, it is reasonable to speculate that the female-specific astrocytic changes in the cortex and hippocampus (albeit unclear why this would not involve striatum or hypothalamus) could contribute to the accelerated recovery of female hamsters, perhaps through behavioural modification.

## Conclusions

In summary, our data indicate a transient and sex-specific increase in GFAP distribution in the female cortex of hamsters in response to mild COVID-19 without changes in overall astrocyte numbers. In addition to providing the first spatiotemporal insight into astrogliosis in mild respiratory COVID-19, our findings contribute further to the literature suggesting astrocytes in males and females respond differently to inflammatory stimuli, and to recent findings in human patients suggesting astrocyte reactivity, at least in terms of GFAP distribution, may be protective to whole-body health.

## List of abbreviations

ACE2: angiotensin-converting enzyme 2
COVID-19: coronavirus disease 2019
ddH_2_O: double distilled water
DPBS: Dulbecco’s phosphate-buffered saline
dpi: days post-inoculation
GFAP: glial fibrillary acidic protein
GO: Gene Ontology
NEUN: neuronal nuclei
ROI: region of interest
SARS-CoV-2: severe acute respiratory syndrome coronavirus 2
SEM: scanning electron microscopy
SOX9: SRY-box transcription factor 9
TCID50: 50% tissue culture infectious dose

## Ethics approval and consent to participate

All animal work was conducted in compliance with the guidelines established by the Canadian Council on Animal Care, as approved by the Animal Care Committee at the Canadian Science Center for Human and Animal Health (animal use document H-20-006).

## Consent for Publication

Not applicable

## Availability of Data and Materials

These data were previously included in the MSc thesis of MRM ((68); link: https://dspace.library.uvic.ca/items/d7256e07-eee8-4b45-80e7-3c4e4404c299). The MATLAB codes are freely available on GitHub: https://github.com/SwayneLab/Quantitative-analysis-of-astrocyte-properties-in-a-Syrian-hamster-model-of-COVID-19.

The confocal micrographs and pipeline output datasets generated during this study are uploaded under embargo to the Federated Research Data Repository under accession number 10.20383/103.01244. These will become freely available after external review. Link to the metadata and any other information is available from the authors upon reasonable request.

## Authors’ Contributions

LAS, MET and DK conceptualized the project. RV, DK and BW performed the animal inoculations and tissue collection. MRM, LWS, HAV, and LRP processed the tissue. MRM and LWS performed immunostaining and imaging experiments. MRM designed the quantitative pipeline and analyzed images. MRM and NY managed the data and made the first draft pipeline output plots. HAV, LRP and MK performed scanning electron microscopy experiments. MRM, LWS, HAV, and LB made the Figures. These data and some Figures are previously included in the MSc thesis of MRM ((68); link: https://dspace.library.uvic.ca/items/d7256e07-eee8-4b45-80e7-3c4e4404c299). LWS and LAS wrote the manuscript based on the methods and data from the MSc thesis of MRM. All authors read and approved of the final manuscript.

## Acknowledgements

Not applicable.

