## Supplemental Materials for "Transient, early, female-specific increase in cortical glial fibrillary acidic protein distribution in the Syrian hamster model of mild peripheral COVID-19"

#### Development of our novel unbiased quantification pipeline: thresholding

Thresholding, or binarization, was a key component in the development of our quantification pipeline. This step segmented our confocal images into signal and noise: pixels were assigned binary values based on whether their intensity exceeded a specific threshold value (ʈ) ((1,2); reviewed in (3)). We compared four thresholding options in the development of our pipeline: Otsu Method, Kapur's Entropy Thresholding, Moment-Preserving Thresholding Method, and Minimum Error Thresholding Method (See Figure S1A and Table S2). Note that given our tiled confocal datasets showed consistent signal to noise across entire images, we only included global thresholding techniques in our comparison (4).

We calculated the peak signal-to-noise ratio (PSNR) to quantitatively evaluate the quality of each thresholding technique on our confocal dataset. This metric compares thresholded images against a predefined ground truth. The PSNR formula is expressed as:

$$\mathrm{PSNR} = 20 \cdot \log_{10}\left(\frac{\mathrm{MAX}_I}{\sqrt{\mathrm{MSE}}}\right)$$

Ground truth (the ideal result) was established by manual adjustment of the threshold levels for three representative images (Figure S1A). Using these ground truth images, we calculated the mean squared error (MSE) between the ground truth and the images produced by each thresholding method. MSE is defined as:

$$\text{MSE} = \frac{1}{MN}\sum_{i=1}^{M}\sum_{j=1}^{N}\big(I(i,j) - K(i,j)\big)^2$$

Where, I(i,j) represents the pixel value in the ground truth image, K(i,j) is the pixel value in the experimental image, and M and N denote the dimensions of the images.

We calculated the PSNR (reported in decibels; dB) using the MSE, the maximum possible pixel value ($MAX_I$; 255 for an 8-bit image), and the peak signal value. The results are shown in Figure S1B and Table S3, with higher PSNRs indicating better thresholding performance (more closely matched to ground truth images). In each example image, Kapur's entropy thresholding yielded the highest PSNR. As such, we included this thresholding technique in the final form of our quantitative analysis pipeline.

### Supplementary Tables

**Table S1. Statistical outcomes from weight tracking data of infected Syrian hamsters** (Figure 1C). Hamsters were weighed at each day post-inoculation (dpi) up to 20 dpi. Data are presented as % initial weight. Mixed-effect analysis (due to removal of some animals at 1-, 3-, 5-, and 7- dpi endpoints: decreasing sample size (N)) indicated significance of Time ($p < 0.0001$), Sex ($p < 0.0001$), and the Time x Sex interaction ($p < 0.0001$). Post-hoc Šídák's multiple comparisons test outcomes for males *vs*. females at each dpi are listed below.

| dpi | Mean Diff. | 95.00% CI of diff. | Adjusted P Value | Sample size (N) |
|---|---|---|---|---|
| **0** | 0 | -- | -- | 25 |
| **1** | -1.65 | -2.58 to -0.73 | 0.0008 (***) | |
| **2** | -1.08 | -2.08 to -0.066 | 0.0374 (*) | 20 |
| **3** | -2.27 | -4.47 to -0.074 | 0.0433 (*) | |
| **4** | -2.33 | -5.68 to 1.01 | 0.1611 | 15 |
| **5** | -2.33 | -6.36 to 1.71 | 0.2417 | |
| **6** | -4.64 | -11.52 to 2.24 | 0.1647 | 10 |
| **7** | -7.33 | -15.34 to 0.68 | 0.0691 | |
| **8** | -11.82 | -21.07 to -2.57 | 0.0225 (*) | 5 |
| **9** | -11.40 | -19.26 to -3.54 | 0.0142 (*) | |
| **10** | -11.30 | -18.25 to -4.35 | 0.0093 (**) | |
| **11** | -13.48 | -20.50 to -6.46 | 0.0041 (**) | |
| **12** | -11.64 | -18.06 to -5.22 | 0.0047 (**) | |
| **13** | -11.40 | -17.59 to -5.21 | 0.0038 (**) | |
| **14** | -12.16 | -17.90 to -6.42 | 0.0016 (**) | |
| **15** | -15.14 | -21.59 to -8.69 | 0.0009 (***) | |
| **16** | -15.32 | -21.39 to -9.25 | 0.0007 (***) | |
| **17** | -13.04 | -19.33 to -6.75 | 0.0016 (**) | |
| **18** | -14.26 | -20.64 to -7.89 | 0.0010 (***) | |
| **19** | -16.00 | -23.39 to -8.61 | 0.0012 (**) | |
| **20** | -14.22 | -21.51 to -6.93 | 0.0020 (**) | |

**Table S2. Summary of the key characteristics and main approaches of the included thresholding techniques.**

| Thresholding Technique | Key Characteristics | Main Approach | References |
|---|---|---|---|
| Otsu Method | Minimizes within-class variance and maximizes between-class variance to enhance contrast between objects and background | Classifies pixels into two groups (darker or lighter than threshold) and finds the threshold that minimizes the sum of the within-class variances for these groups. It computes the optimal threshold by minimizing the function representing the weighted sum of variances. | (5) |
| Minimum Error Thresholding | Treats the gray-level histogram as a probability density function assuming two normally distributed mixed populations | Sets a criterion function based on parameters of the mixed distribution model to minimize the probability of misclassification of pixels. The optimal threshold maximizes the likelihood of correct classification by minimizing the criterion function involving the standard deviations and probabilities of the distributions separated by the threshold. | (6) |
| Moment-Preserving Thresholding | Calculates threshold values to preserve the statistical moments of the original image's gray-level distribution in the resultant binary image | Threshold values are derived from the image histogram using coefficients related to the moments of the gray level distribution. The process involves calculating moments from pixel intensities and frequencies, and the threshold is determined using a formula that incorporates these moments to maintain the underlying statistical properties of the image. | (4) |
| Kapur's Entropy Thresholding | Uses entropy of the histogram to maximize the information content between segmented parts of an image, based on information theory principles | Initially defines the probability distributions for image thresholding at each potential threshold, splitting the histogram into two segments. The entropy for each segment is calculated to measure the randomness or unpredictability within those segments. The combined entropy for each threshold is computed, and the optimal threshold is the one that maximizes the sum of the entropies of the two segments, reflecting maximum information separation. | (2,7) |

**Table S3. Calculated PSNR values for example images** (Figure S1). Higher values indicate the thresholding method more closely matched ground truth images.

| | *Otsu* | *Kapur* | *Moment* | *Minimum Error* |
|---|---|---|---|---|
| *Example 1* | 2.93 | 19.82 | 2.10 | 10.86 |
| *Example 2* | 6.86 | 7.73 | 2.04 | 6.44 |

| | | | | |
|---|---|---|---|---|
| *Example 3* | 2.85 | 19.59 | 2.22 | 9.35 |

**Table S4. Statistical outcomes for GFAP distribution data** (Figure 4C,D and S3). GFAP distribution data from uninfected controls (CTL) and each experimental group (1-, 3-, 5-, 7-, 31-dpi) were analyzed by one-way ANOVA with post-hoc Dunnet's test (all compared against CTL) for each brain ROI. ANOVA results are shown in rows labelled "ANOVA (Overall)", and all other rows correspond to the post-hoc test results. Lowerdiff: The lower bound of the 95% confidence interval for the estimated difference between the experimental condition and the control. ESTdiff: The point estimate of the difference between the experimental condition and the control. Upperdiff: The upper bound of the 95% confidence interval for the estimated difference.

| ROI | Sex | Comparison | Lowerdiff | ESTdiff | Upperdiff | p | F(2,14) | $\eta^2$ |
|---|---|---|---|---|---|---|---|---|
| Cortex | F | **ANOVA (Overall)** | | | | **0.00694** | **5.3396** | **0.6725** |
| | | 1 vs. CTL | -24.20 | 52.81 | 129.82 | 0.2323 | | |
| | | **3 vs. CTL** | **7.18** | **79.22** | **151.25** | **0.0297** | | |
| | | 5 vs. CTL | -93.13 | -16.12 | 60.89 | 0.9581 | | |
| | | 7 vs. CTL | -97.96 | -20.95 | 56.06 | 0.8920 | | |
| | | 31 vs. CTL | -80.28 | -3.27 | 73.74 | 1.0000 | | |
| | M | ANOVA (Overall) | | | | 0.4391 | 1.0321 | 0.2842 |
| | | 1 vs. CTL | -77.85 | 35.97 | 149.79 | 0.8271 | | |
| | | 3 vs. CTL | -135.60 | -21.78 | 92.04 | 0.9706 | | |
| | | 5 vs. CTL | -54.65 | 51.82 | 158.29 | 0.5162 | | |
| | | 7 vs. CTL | -74.76 | 39.06 | 152.88 | 0.7812 | | |
| | | 31 vs. CTL | -86.80 | 27.02 | 140.84 | 0.9329 | | |
| Corpus callosum | F | **ANOVA (Overall)** | | | | **0.0316** | **3.4181** | **0.5497** |
| | | 1 vs. CTL | -139.19 | -32.47 | 74.25 | 0.8485 | | |
| | | 3 vs. CTL | -15.49 | 79.97 | 175.42 | 0.1147 | | |
| | | 5 vs. CTL | -100.51 | 6.22 | 112.94 | 0.9999 | | |
| | | 7 vs. CTL | -140.67 | -33.94 | 72.78 | 0.8268 | | |
| | | 31 vs. CTL | -109.14 | -2.41 | 104.31 | 1.0000 | | |
| | M | ANOVA (Overall) | | | | 0.3194 | 1.3207 | 0.3550 |
| | | 1 vs. CTL | -135.39 | 67.70 | 270.80 | 0.7945 | | |
| | | 3 vs. CTL | -229.02 | -25.92 | 177.17 | 0.9947 | | |
| | | 5 vs. CTL | -81.34 | 121.75 | 324.85 | 0.3323 | | |
| | | 7 vs. CTL | -110.61 | 92.49 | 295.58 | 0.5681 | | |
| | | 31 vs. CTL | -123.84 | 79.25 | 282.34 | 0.6909 | | |
| | | ANOVA (Overall) | | | | 0.4061 | 1.1078 | 0.3158 |

| | | | | | | | |
|---|---|---|---|---|---|---|---|
| Hippocampus | F | 1 vs. CTL | -94.20 | 174.04 | 442.29 | 0.2697 | | |
| | | 3 vs. CTL | -80.85 | 187.39 | 455.64 | 0.2164 | | |
| | | 5 vs. CTL | -116.25 | 151.99 | 420.24 | 0.3794 | | |
| | | 7 vs. CTL | -176.11 | 92.13 | 360.38 | 0.7767 | | |
| | | 31 vs. CTL | -151.13 | 117.12 | 385.36 | 0.6031 | | |
| | M | ANOVA (Overall) | | | | 0.4011 | 1.1112 | 0.2994 |
| | | 1 vs. CTL | -91.86 | 38.57 | 169.00 | 0.8588 | | |
| | | 3 vs. CTL | -180.97 | -50.54 | 79.89 | 0.7005 | | |
| | | 5 vs. CTL | -150.80 | -20.38 | 110.05 | 0.9874 | | |
| | | 7 vs. CTL | -99.02 | 31.41 | 161.83 | 0.9293 | | |
| | | 31 vs. CTL | -104.58 | 17.43 | 139.43 | 0.9915 | | |
| Third ventricle | F | ANOVA (Overall) | | | | 0.7175 | 0.5765 | 0.1413 |
| | | 1 vs. CTL | -145.90 | -32.29 | 81.32 | 0.8754 | | |
| | | 3 vs. CTL | -125.22 | -11.61 | 102.00 | 0.9981 | | |
| | | 5 vs. CTL | -119.55 | -13.28 | 92.99 | 0.9953 | | |
| | | 7 vs. CTL | -119.62 | -6.01 | 107.59 | 0.9999 | | |
| | | 31 vs. CTL | -170.94 | -57.33 | 56.28 | 0.4844 | | |
| | M | ANOVA (Overall) | | | | 0.8935 | 0.3168 | 0.0811 |
| | | 1 vs. CTL | -146.98 | 10.59 | 168.17 | 0.9997 | | |
| | | 3 vs. CTL | -196.51 | -38.94 | 118.63 | 0.9206 | | |
| | | 5 vs. CTL | -134.00 | 23.58 | 181.15 | 0.9892 | | |
| | | 7 vs. CTL | -172.14 | -14.57 | 143.00 | 0.9988 | | |
| | | 31 vs. CTL | -165.54 | -7.97 | 149.60 | 0.9999 | | |
| Striatum | F | ANOVA (Overall) | | | | 0.8430 | 0.395 | 0.1815 |
| | | 1 vs. CTL | -164.77 | -38.62 | 87.53 | 0.8397 | | |
| | | 3 vs. CTL | -156.10 | -29.95 | 96.20 | 0.9311 | | |
| | | 5 vs. CTL | -176.37 | -50.22 | 75.93 | 0.6761 | | |
| | | 7 vs. CTL | -150.68 | -24.53 | 101.63 | 0.9678 | | |
| | | 31 vs. CTL | -178.40 | -52.25 | 73.90 | 0.6457 | | |
| | M | ANOVA (Overall) | | | | 0.8935 | 0.2118 | 0.1166 |
| | | 1 vs. CTL | -146.97 | 10.59 | 168.15 | 0.9997 | | |
| | | 3 vs. CTL | -196.50 | -38.94 | 118.62 | 0.9206 | | |
| | | 5 vs. CTL | -133.98 | 23.58 | 181.14 | 0.9892 | | |
| | | 7 vs. CTL | -172.13 | -14.57 | 142.99 | 0.9988 | | |
| | | 31 vs. CTL | -165.53 | -7.97 | 149.59 | 0.9999 | | |

**Table S5. Statistical outcomes for SOX9-positive cell density data** (Figure S4). SOX9-positive cell density data from uninfected controls (CTL) and each experimental group (1-, 3-, 5-, 7-, 31-dpi) were analyzed by one-way ANOVA with post-hoc Dunnet's test (all compared against CTL) for each brain ROI. ANOVA results are shown in rows labelled "ANOVA (Overall)", and all other rows correspond to the post-hoc test results. Lowerdiff: The lower bound of the 95% confidence interval for the estimated difference between the experimental condition and the control. ESTdiff: The point estimate of the difference between the experimental condition and the control. Upperdiff: The upper bound of the 95% confidence interval for the estimated difference.

| ROI | Sex | Comparison | Lowerdiff | ESTdiff | Upperdiff | p | F(2,14) | $\eta^2$ |
|---|---|---|---|---|---|---|---|---|
| Cortex | F | ANOVA (Overall) | | | | 0.7731 | 0.4966 | 0.1714 |
| | | 1 vs. CTL | -70.59 | 49.25 | 169.09 | 0.6518 | | |
| | | 3 vs. CTL | -115.60 | 4.24 | 124.08 | 1.0000 | | |
| | | 5 vs. CTL | -117.26 | 2.59 | 122.43 | 1.0000 | | |
| | | 7 vs. CTL | -82.21 | 37.63 | 157.47 | 0.8271 | | |
| | | 31 vs. CTL | -104.37 | 15.47 | 135.31 | 0.9944 | | |
| | M | ANOVA (Overall) | | | | 0.0702 | 2.7498 | 0.5340 |
| | | 1 vs. CTL | -166.24 | 34.33 | 234.90 | 0.9809 | | |
| | | 3 vs. CTL | -39.92 | 160.65 | 361.22 | 0.1344 | | |
| | | 5 vs. CTL | -72.35 | 128.22 | 328.79 | 0.2811 | | |
| | | 7 vs. CTL | -206.54 | -5.97 | 194.60 | 1.0000 | | |
| | | 31 vs. CTL | -30.22 | 170.34 | 370.91 | 0.1064 | | |
| Corpus callosum | F | ANOVA (Overall) | | | | 0.8887 | 0.3244 | 0.1191 |
| | | 1 vs. CTL | -100.27 | -3.03 | 94.21 | 1.0000 | | |
| | | 3 vs. CTL | -90.78 | 6.47 | 103.71 | 0.9998 | | |
| | | 5 vs. CTL | -123.96 | -26.72 | 70.53 | 0.8861 | | |
| | | 7 vs. CTL | -99.67 | -2.43 | 94.81 | 1.0000 | | |
| | | 31 vs. CTL | -120.47 | -23.23 | 74.02 | 0.9297 | | |
| | M | ANOVA (Overall) | | | | 0.9794 | 0.1401 | 0.0551 |
| | | 1 vs. CTL | -147.27 | -23.15 | 100.97 | 0.9728 | | |
| | | 3 vs. CTL | -117.11 | 7.01 | 131.13 | 0.9999 | | |
| | | 5 vs. CTL | -118.18 | 5.94 | 130.06 | 1.0000 | | |
| | | 7 vs. CTL | -122.04 | 2.08 | 126.20 | 1.0000 | | |
| | | 31 vs. CTL | -131.70 | -7.58 | 116.54 | 0.9998 | | |
| Hippocampus | F | ANOVA (Overall) | | | | 0.7955 | 0.4645 | 0.1622 |
| | | 1 vs. CTL | -140.90 | 121.99 | 384.88 | 0.5522 | | |
| | | 3 vs. CTL | -218.17 | 44.72 | 307.61 | 0.9813 | | |
| | | 5 vs. CTL | -224.54 | 38.35 | 301.24 | 0.9903 | | |
| | | 7 vs. CTL | -240.25 | 22.64 | 285.53 | 0.9991 | | |
| | | 31 vs. CTL | -252.66 | 10.23 | 273.12 | 1.0000 | | |
| | M | ANOVA (Overall) | | | | 0.5553 | 0.8225 | 0.2403 |
| | | 1 vs. CTL | -108.69 | -12.77 | 83.14 | 0.9938 | | |
| | | 3 vs. CTL | -125.89 | -29.97 | 65.94 | 0.8328 | | |
| | | 5 vs. CTL | -114.22 | -18.31 | 77.61 | 0.9709 | | |
| | | 7 vs. CTL | -157.86 | -61.94 | 33.97 | 0.2768 | | |
| | | 31 vs. CTL | -124.13 | -34.41 | 55.31 | 0.7078 | | |
| Third ventricle | F | ANOVA (Overall) | | | | 0.0640 | 2.7777 | 0.5165 |
| | | 1 vs. CTL | -49.87 | 31.18 | 112.24 | 0.7055 | | |
| | | 3 vs. CTL | -19.13 | 61.92 | 142.97 | 0.1621 | | |
| | | 5 vs. CTL | -92.87 | -17.05 | 58.76 | 0.9450 | | |
| | | 7 vs. CTL | -28.67 | 52.38 | 133.43 | 0.2762 | | |
| | | 31 vs. CTL | -34.77 | 46.28 | 127.33 | 0.3772 | | |
| | M | ANOVA (Overall) | | | | 0.1028 | 2.3208 | 0.4716 |
| | | 1 vs. CTL | -57.79 | 6.77 | 71.33 | 0.9979 | | |
| | | 3 vs. CTL | -94.89 | -30.34 | 34.22 | 0.5464 | | |
| | | 5 vs. CTL | -85.78 | -21.22 | 43.34 | 0.8061 | | |
| | | 7 vs. CTL | -118.69 | -54.14 | 10.42 | 0.1132 | | |

| | | | | | | | | |
|---|---|---|---|---|---|---|---|---|
| | | 31 vs. CTL | -57.06 | 3.33 | 63.71 | 0.9999 | | |
| Striatum | F | ANOVA (Overall) | | | | 0.2335 | 1.5810 | 0.3781 |
| | | 1 vs. CTL | -148.98 | -14.82 | 119.34 | 0.9973 | | |
| | | 3 vs. CTL | -195.49 | -61.33 | 72.83 | 0.5697 | | |
| | | 5 vs. CTL | -203.50 | -78.00 | 47.49 | 0.3069 | | |
| | | 7 vs. CTL | -227.16 | -93.00 | 41.16 | 0.2246 | | |
| | | 31 vs. CTL | -137.49 | -3.33 | 130.83 | 1.0000 | | |
| | M | ANOVA (Overall) | | | | 0.1028 | 0.3675 | 0.1328 |
| | | 1 vs. CTL | -57.78 | 6.77 | 71.32 | 0.9979 | | |
| | | 3 vs. CTL | -94.89 | -30.34 | 34.22 | 0.5464 | | |
| | | 5 vs. CTL | -85.77 | -21.22 | 43.33 | 0.8061 | | |
| | | 7 vs. CTL | -118.69 | -54.14 | 10.42 | 0.1131 | | |
| | | 31 vs. CTL | -57.06 | 3.33 | 63.71 | 0.9999 | | |

**Table S6. Statistical outcomes for NEUN-positive cell density data** (Figure S5). NEUN-positive cell density data from uninfected controls (CTL) and each experimental group (1-, 3-, 5-, 7-, 31-dpi) were analyzed by one way ANOVA with post-hoc Dunnet's test (all compared against CTL) for each brain ROI. ANOVA results are shown in rows labelled "ANOVA (Overall)", and all other rows correspond to the post-hoc test results. Lowerdiff: The lower bound of the 95% confidence interval for the estimated difference between the experimental condition and the control. ESTdiff: The point estimate of the difference between the experimental condition and the control. Upperdiff: The upper bound of the 95% confidence interval for the estimated difference.

| ROI | Sex | Comparison | Lowerdiff | ESTdiff | Upperdiff | p | $F(2,14)$ | $\eta^2$ |
|---|---|---|---|---|---|---|---|---|
| Cortex | F | ANOVA (Overall) | | | | 0.2517 | 1.4675 | 0.3015 |
| | | 1 vs. CTL | -125.09 | -21.73 | 81.62 | 0.9691 | | |
| | | 3 vs. CTL | -146.22 | -56.71 | 32.79 | 0.3186 | | |
| | | 5 vs. CTL | -170.39 | -75.45 | 19.49 | 0.1497 | | |
| | | 7 vs. CTL | -142.43 | -39.07 | 64.28 | 0.7606 | | |
| | | 31 vs. CTL | -106.27 | -2.91 | 100.44 | 1.0000 | | |
| | M | ANOVA (Overall) | | | | 0.6557 | 0.6663 | 0.2040 |
| | | 1 vs. CTL | -120.05 | 12.06 | 144.18 | 0.9989 | | |
| | | 3 vs. CTL | -195.38 | -63.26 | 68.85 | 0.5303 | | |
| | | 5 vs. CTL | -162.55 | -30.43 | 101.68 | 0.9399 | | |
| | | 7 vs. CTL | -151.78 | -19.66 | 112.45 | 0.9898 | | |
| | | 31 vs. CTL | -155.45 | -31.87 | 91.71 | 0.9100 | | |
| Corpus | F | ANOVA (Overall) | | | | 0.4461 | 1.0090 | 0.2517 |
| | | 1 vs. CTL | -167.34 | -48.30 | 70.75 | 0.6716 | | |

| | | | | | | | | |
|---|---|---|---|---|---|---|---|---|
| callosum | | 3 vs. CTL | -201.75 | -82.70 | 36.35 | 0.2272 | | |
| | | 5 vs. CTL | -199.90 | -80.85 | 38.19 | 0.2434 | | |
| | | 7 vs. CTL | -200.09 | -72.82 | 54.45 | 0.3812 | | |
| | | 31 vs. CTL | -189.37 | -62.10 | 65.17 | 0.5198 | | |
| | M | ANOVA (Overall) | | | | 0.1116 | 2.2874 | 0.4880 |
| | | 1 vs. CTL | -85.94 | 9.16 | 104.25 | 0.9985 | | |
| | | 3 vs. CTL | -163.35 | -68.25 | 26.84 | 0.1985 | | |
| | | 5 vs. CTL | -144.90 | -49.81 | 45.29 | 0.4474 | | |
| | | 7 vs. CTL | -90.02 | 5.08 | 100.17 | 0.9999 | | |
| | | 31 vs. CTL | -151.40 | -56.30 | 38.79 | 0.3427 | | |
| Hippocampus | F | ANOVA (Overall) | | | | 0.2920 | 1.3864 | 0.3478 |
| | | 1 vs. CTL | -120.65 | 64.66 | 249.97 | 0.7710 | | |
| | | 3 vs. CTL | -188.52 | -3.20 | 182.11 | 1.0000 | | |
| | | 5 vs. CTL | -244.97 | -71.62 | 101.72 | 0.6522 | | |
| | | 7 vs. CTL | -205.00 | -19.69 | 165.63 | 0.9978 | | |
| | | 31 vs. CTL | -129.53 | 55.78 | 241.09 | 0.8507 | | |
| | M | ANOVA (Overall) | | | | 0.1069 | 2.2846 | 0.4677 |
| | | 1 vs. CTL | -126.56 | 24.99 | 176.55 | 0.9841 | | |
| | | 3 vs. CTL | -210.12 | -58.56 | 93.00 | 0.7024 | | |
| | | 5 vs. CTL | -78.88 | 72.68 | 224.23 | 0.5288 | | |
| | | 7 vs. CTL | -200.25 | -48.69 | 102.87 | 0.8184 | | |
| | | 31 vs. CTL | -207.76 | -65.99 | 75.78 | 0.5542 | | |
| Third ventricle | F | ANOVA (Overall) | | | | 0.6556 | 0.6664 | 0.2040 |
| | | 1 vs. CTL | -141.25 | -1.85 | 137.54 | 1.0000 | | |
| | | 3 vs. CTL | -133.93 | 5.47 | 144.86 | 1.0000 | | |
| | | 5 vs. CTL | -173.66 | -43.27 | 87.12 | 0.8007 | | |
| | | 7 vs. CTL | -101.10 | 38.29 | 177.68 | 0.8886 | | |
| | | 31 vs. CTL | -144.27 | -4.88 | 134.52 | 1.0000 | | |
| | M | ANOVA (Overall) | | | | 0.5856 | 0.7756 | 0.2442 |
| | | 1 vs. CTL | -150.37 | -29.25 | 91.88 | 0.9269 | | |
| | | 3 vs. CTL | -170.30 | -49.17 | 71.95 | 0.6612 | | |
| | | 5 vs. CTL | -99.64 | 21.48 | 142.61 | 0.9779 | | |
| | | 7 vs. CTL | -150.70 | -29.57 | 91.55 | 0.9239 | | |
| | | 31 vs. CTL | -120.51 | 0.61 | 121.74 | 1.0000 | | |
| Striatum | F | ANOVA (Overall) | | | | 0.6316 | 0.7024 | 0.2127 |
| | | 1 vs. CTL | -300.59 | 148.96 | 598.51 | 0.8016 | | |
| | | 3 vs. CTL | -401.11 | 48.44 | 497.99 | 0.9976 | | |
| | | 5 vs. CTL | -531.57 | -111.05 | 309.47 | 0.9023 | | |
| | | 7 vs. CTL | -491.74 | -42.19 | 407.36 | 0.9988 | | |
| | | 31 vs. CTL | -421.97 | 27.58 | 477.13 | 0.9998 | | |
| | M | ANOVA (Overall) | | | | 0.4269 | 1.0632 | 0.3070 |
| | | 1 vs. CTL | -256.62 | -13.58 | 229.47 | 0.9999 | | |
| | | 3 vs. CTL | -231.80 | 11.24 | 254.29 | 1.0000 | | |
| | | 5 vs. CTL | -122.60 | 120.44 | 363.49 | 0.4956 | | |
| | | 7 vs. CTL | -254.62 | -11.57 | 231.47 | 1.0000 | | |
| | | 31 vs. CTL | -306.42 | -63.38 | 179.67 | 0.9044 | | |

**Table S7. Statistical outcomes for Hoechst pixel density data** (Figure S6). Hoechst pixel density data from uninfected controls (CTL) and each experimental group (1-, 3-, 5-, 7-, 31-

dpi) were analyzed by one way ANOVA with post-hoc Dunnet's test (all compared against CTL) for each brain ROI. ANOVA results are shown in rows labelled "ANOVA (Overall)", and all other rows correspond to the post-hoc test results. Lowerdiff: The lower bound of the 95% confidence interval for the estimated difference between the experimental condition and the control. ESTdiff: The point estimate of the difference between the experimental condition and the control. Upperdiff: The upper bound of the 95% confidence interval for the estimated difference.

| ROI | Sex | Comparison | Lowerdiff | ESTdiff | Upperdiff | p | F(2,14) | $\eta^2$ |
|---|---|---|---|---|---|---|---|---|
| Cortex | F | ANOVA (Overall) | | | | 0.8226 | 0.4260 | 0.1408 |
| | | 1 vs. CTL | -47.35 | -1.74 | 43.87 | 1.0000 | | |
| | | 3 vs. CTL | -30.95 | 14.66 | 60.28 | 0.8182 | | |
| | | 5 vs. CTL | -35.57 | 7.09 | 49.76 | 0.9836 | | |
| | | 7 vs. CTL | -41.90 | 3.72 | 49.33 | 0.9994 | | |
| | | 31 vs. CTL | -29.86 | 15.75 | 61.37 | 0.7774 | | |
| | M | ANOVA (Overall) | | | | 0.6742 | 0.6397 | 0.2104 |
| | | 1 vs. CTL | -90.19 | -17.42 | 55.34 | 0.9289 | | |
| | | 3 vs. CTL | -66.79 | 5.98 | 78.74 | 0.9993 | | |
| | | 5 vs. CTL | -83.98 | -11.21 | 61.55 | 0.9877 | | |
| | | 7 vs. CTL | -107.23 | -34.46 | 38.30 | 0.5346 | | |
| | | 31 vs. CTL | -81.59 | -8.83 | 63.94 | 0.9958 | | |
| Corpus callosum | F | ANOVA (Overall) | | | | 0.7384 | 0.5470 | 0.1460 |
| | | 1 vs. CTL | -88.38 | -6.73 | 74.92 | 0.9994 | | |
| | | 3 vs. CTL | -49.39 | 28.68 | 106.75 | 0.7444 | | |
| | | 5 vs. CTL | -55.26 | 26.39 | 108.04 | 0.8204 | | |
| | | 7 vs. CTL | -72.92 | 14.37 | 101.66 | 0.9849 | | |
| | | 31 vs. CTL | -68.31 | 18.98 | 106.26 | 0.9534 | | |
| | M | ANOVA (Overall) | | | | 0.9627 | 0.1854 | 0.0717 |
| | | 1 vs. CTL | -199.09 | -14.91 | 169.27 | 0.9994 | | |
| | | 3 vs. CTL | -170.08 | 14.10 | 198.28 | 0.9995 | | |
| | | 5 vs. CTL | -175.70 | 8.48 | 192.66 | 1.0000 | | |
| | | 7 vs. CTL | -140.59 | 43.59 | 227.77 | 0.9319 | | |
| | | 31 vs. CTL | -173.78 | 10.40 | 194.59 | 0.9999 | | |
| Hippocampus | F | ANOVA (Overall) | | | | 0.2533 | 1.5271 | 0.3889 |
| | | 1 vs. CTL | -58.03 | -10.71 | 36.60 | 0.9422 | | |
| | | 3 vs. CTL | -40.94 | 6.37 | 53.68 | 0.9932 | | |
| | | 5 vs. CTL | -44.61 | 2.70 | 50.02 | 0.9999 | | |
| | | 7 vs. CTL | -80.27 | -32.96 | 14.36 | 0.2185 | | |
| | | 31 vs. CTL | -56.92 | -9.61 | 37.70 | 0.9617 | | |
| | M | ANOVA (Overall) | | | | 0.7880 | 0.4753 | 0.1653 |
| | | 1 vs. CTL | -116.54 | -30.49 | 55.56 | 0.7574 | | |
| | | 3 vs. CTL | -97.72 | -11.67 | 74.38 | 0.9930 | | |
| | | 5 vs. CTL | -101.85 | -15.81 | 70.24 | 0.9744 | | |
| | | 7 vs. CTL | -117.96 | -31.91 | 54.13 | 0.7271 | | |

| | | | | | | | |
|---|---|---|---|---|---|---|---|
| | | 31 vs. CTL | -123.83 | -37.78 | 48.27 | 0.5986 | | |
| Third ventricle | F | ANOVA (Overall) | | | | 0.6129 | 0.7324 | 0.2338 |
| | | 1 vs. CTL | -70.85 | 2.13 | 75.11 | 1.0000 | | |
| | | 3 vs. CTL | -37.14 | 35.84 | 108.82 | 0.5034 | | |
| | | 5 vs. CTL | -40.55 | 32.43 | 105.41 | 0.5886 | | |
| | | 7 vs. CTL | -49.67 | 23.30 | 96.28 | 0.8183 | | |
| | | 31 vs. CTL | -60.33 | 12.65 | 85.63 | 0.9798 | | |
| | M | ANOVA (Overall) | | | | 0.3958 | 1.1229 | 0.3016 |
| | | 1 vs. CTL | -60.89 | 19.27 | 99.44 | 0.9298 | | |
| | | 3 vs. CTL | -108.33 | -28.16 | 52.01 | 0.7669 | | |
| | | 5 vs. CTL | -104.59 | -24.42 | 55.74 | 0.8451 | | |
| | | 7 vs. CTL | -89.22 | -9.05 | 71.11 | 0.9971 | | |
| | | 31 vs. CTL | -57.33 | 17.66 | 92.65 | 0.9348 | | |
| Striatum | F | ANOVA (Overall) | | | | 0.3786 | 1.1699 | 0.3277 |
| | | 1 vs. CTL | -94.17 | 0.88 | 95.92 | 1.0000 | | |
| | | 3 vs. CTL | -54.90 | 40.14 | 135.18 | 0.6302 | | |
| | | 5 vs. CTL | -89.16 | 5.88 | 100.92 | 0.9998 | | |
| | | 7 vs. CTL | -128.83 | -33.79 | 61.26 | 0.7554 | | |
| | | 31 vs. CTL | -70.90 | 24.14 | 119.18 | 0.9126 | | |
| | M | ANOVA (Overall) | | | | 0.9908 | 0.0975 | 0.0391 |
| | | 1 vs. CTL | -92.82 | 2.44 | 97.71 | 1.0000 | | |
| | | 3 vs. CTL | -84.46 | 10.81 | 106.07 | 0.9969 | | |
| | | 5 vs. CTL | -85.26 | 10.00 | 105.27 | 0.9978 | | |
| | | 7 vs. CTL | -104.03 | -8.77 | 86.50 | 0.9988 | | |
| | | 31 vs. CTL | -94.86 | 0.41 | 95.68 | 1.0000 | | |

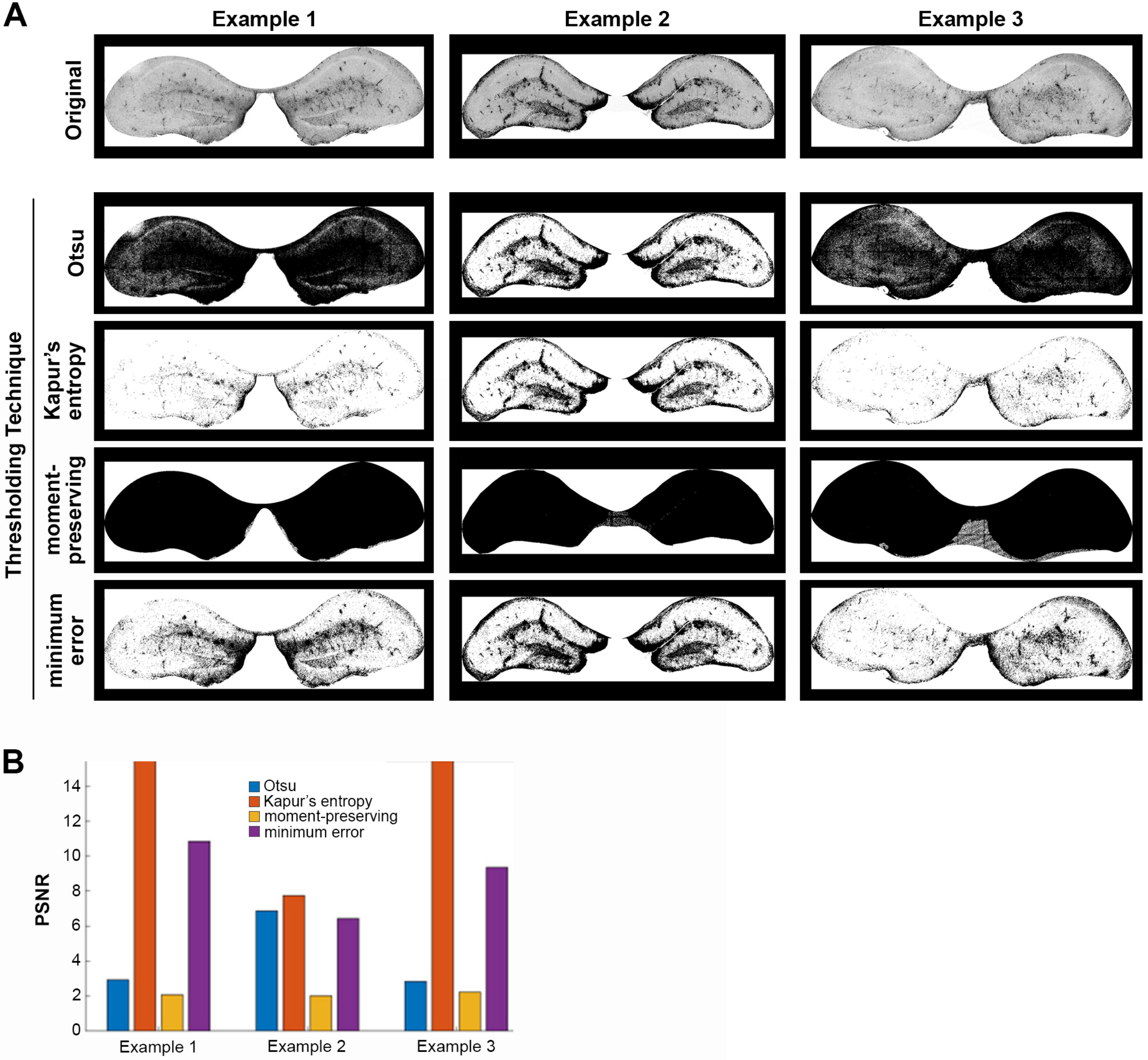
A
Example 1
Example 2
Example 3
Original
Thresholding Technique
Otsu
Kapur's entropy
moment-preserving
minimum error
B
PSNR
Otsu
Kapur's entropy
moment-preserving
minimum error
0
2
4
6
8
10
12
14
Example 1
Example 2
Example 3

Figure S1

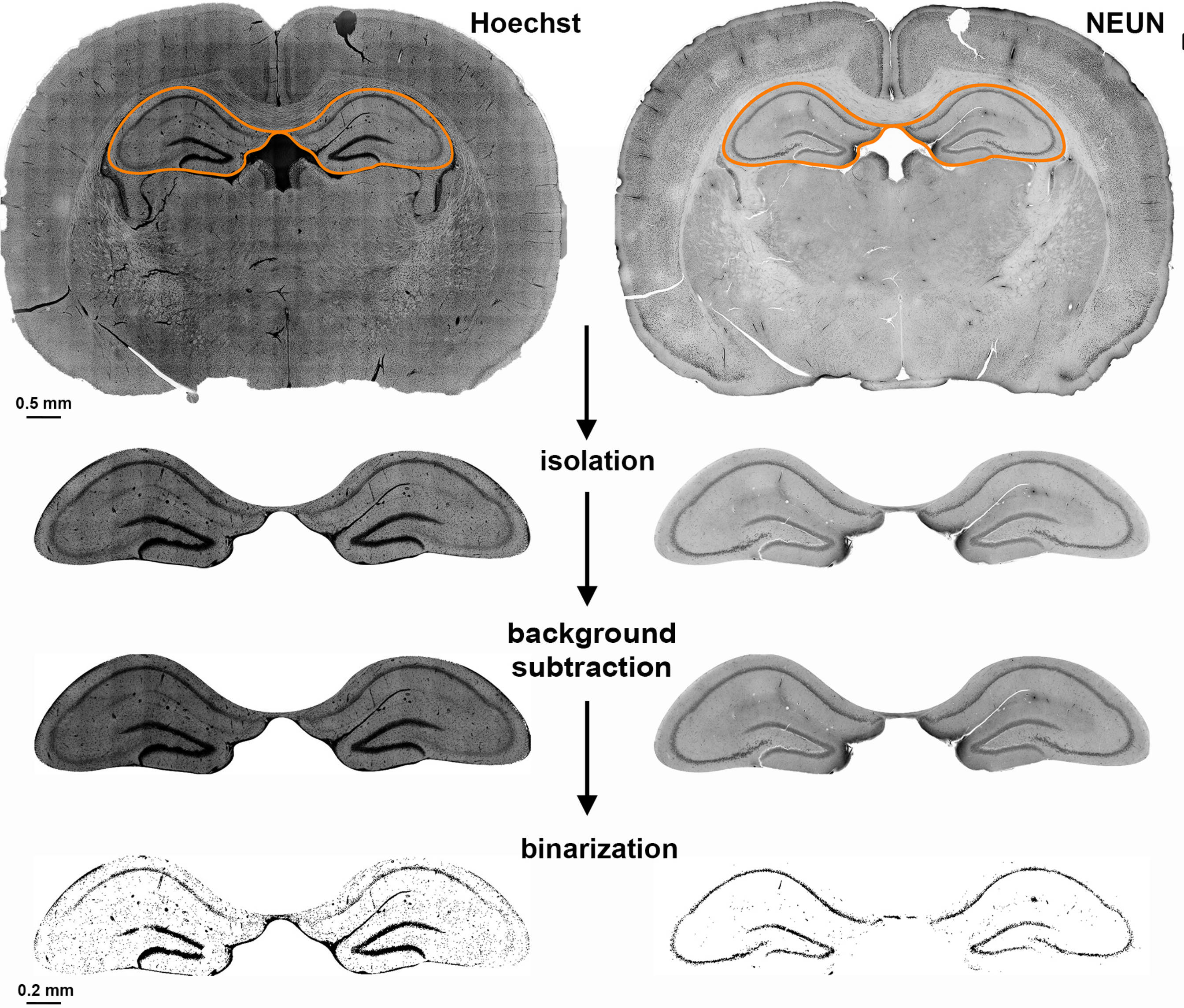


| metrics | value |
|---|---|
| Sum of Hoechst positive pixels | 6.53e+05 |
| Hoechst area (mm$^2$) | 6.53e-01 |
| total area of sample (mm$^2$) | 11.12 |
| Percentage of Hoechst coverage | 5.86% |

| metrics | value |
|---|---|
| NeuN positive cell number | 3.88e+05 |
| NeuN area (mm$^2$) | 0.38 |
| total area of sample (mm$^2$) | 11.12 |
| percentage NeuN coverage | 3.49% |

*Figure S2*

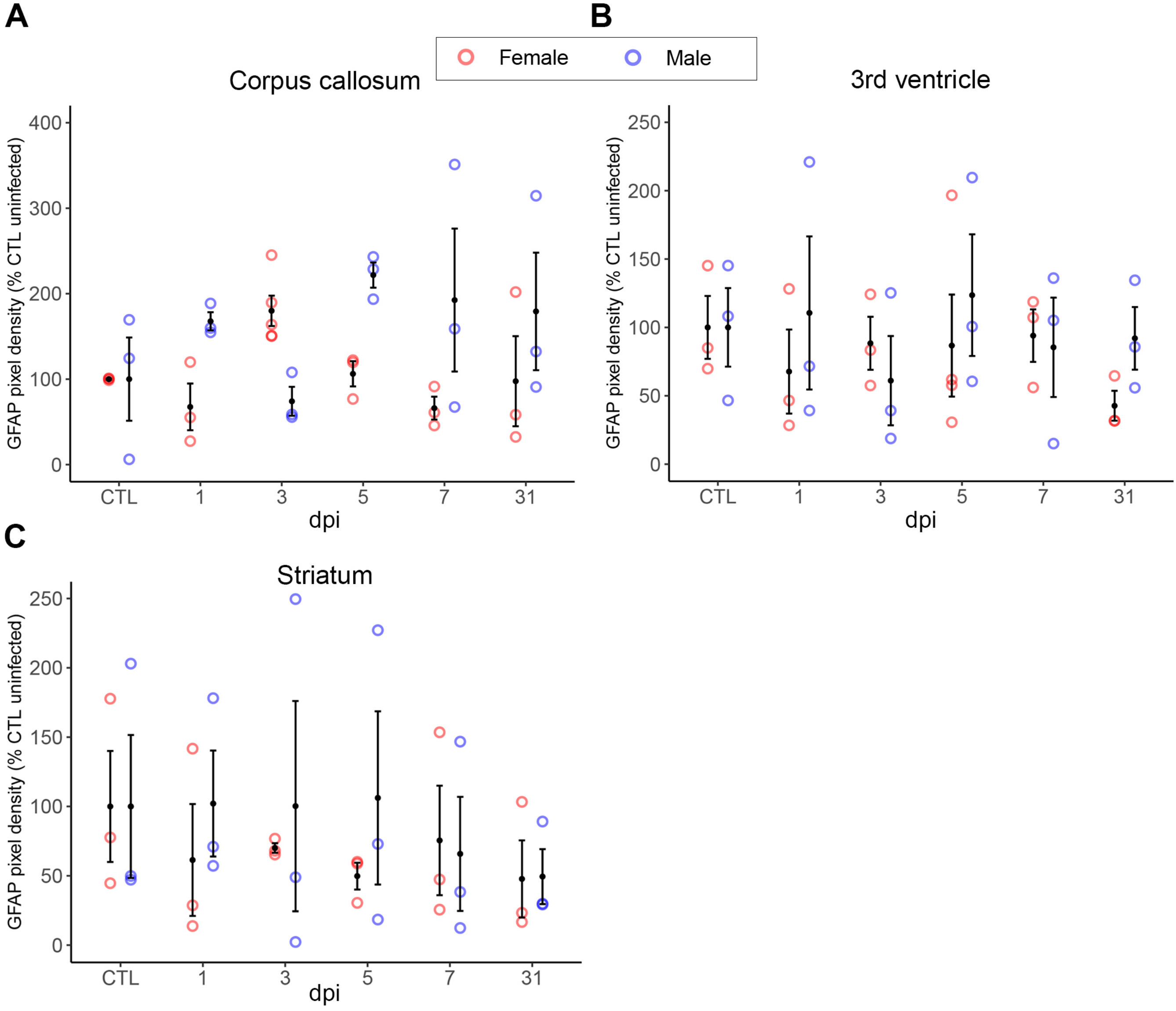
A
B
C
Female
Male
Corpus callosum
3rd ventricle
Striatum
GFAP pixel density (% CTL uninfected)
dpi
CTL
1
3
5
7
31
0
100
200
300
400
50
150
250

Figure S3

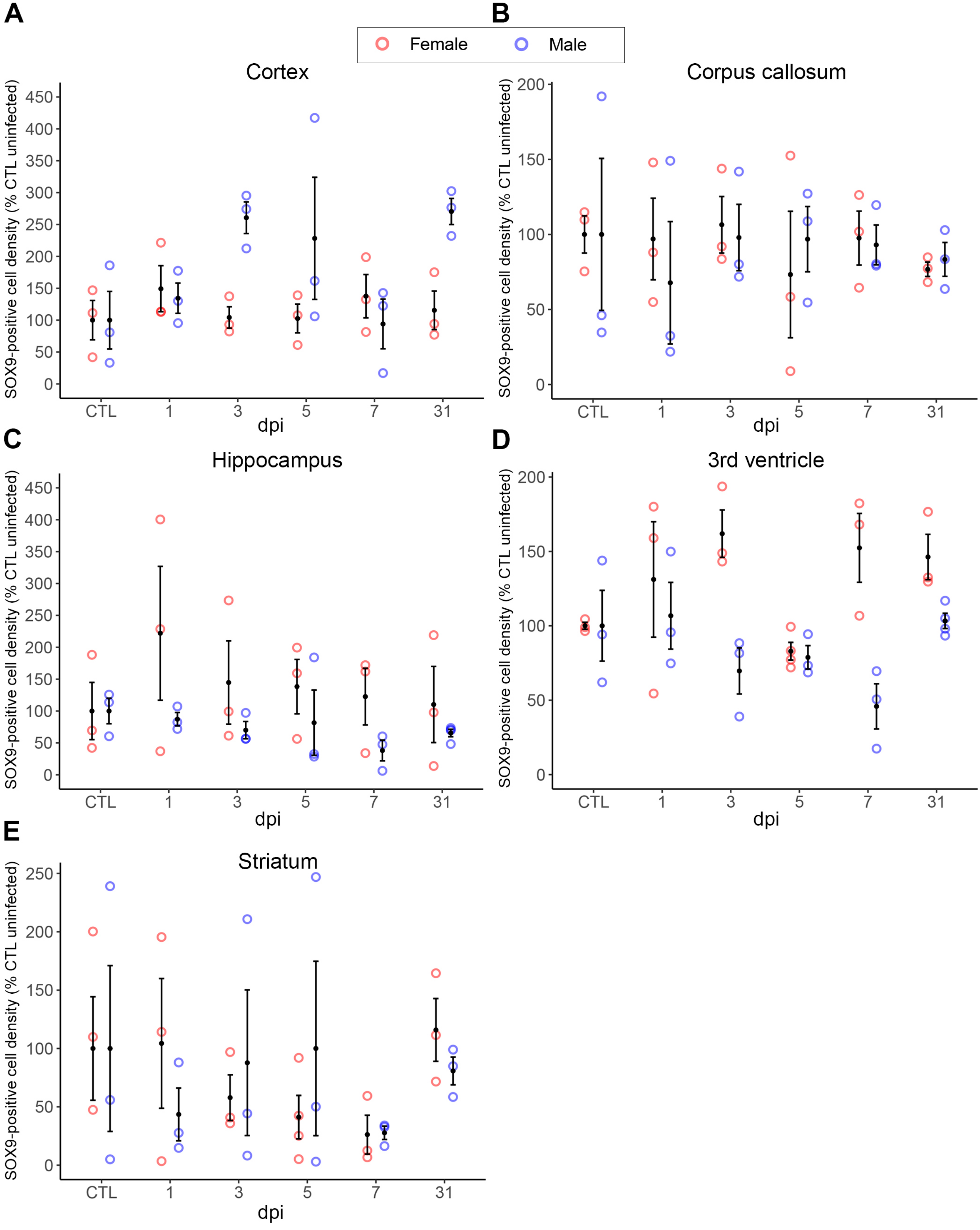
A
B
Female
Male
Cortex
Corpus callosum
SOX9-positive cell density (% CTL uninfected)
dpi
CTL
1
3
5
7
31
C
D
Hippocampus
3rd ventricle
E
Striatum

*Figure S4*

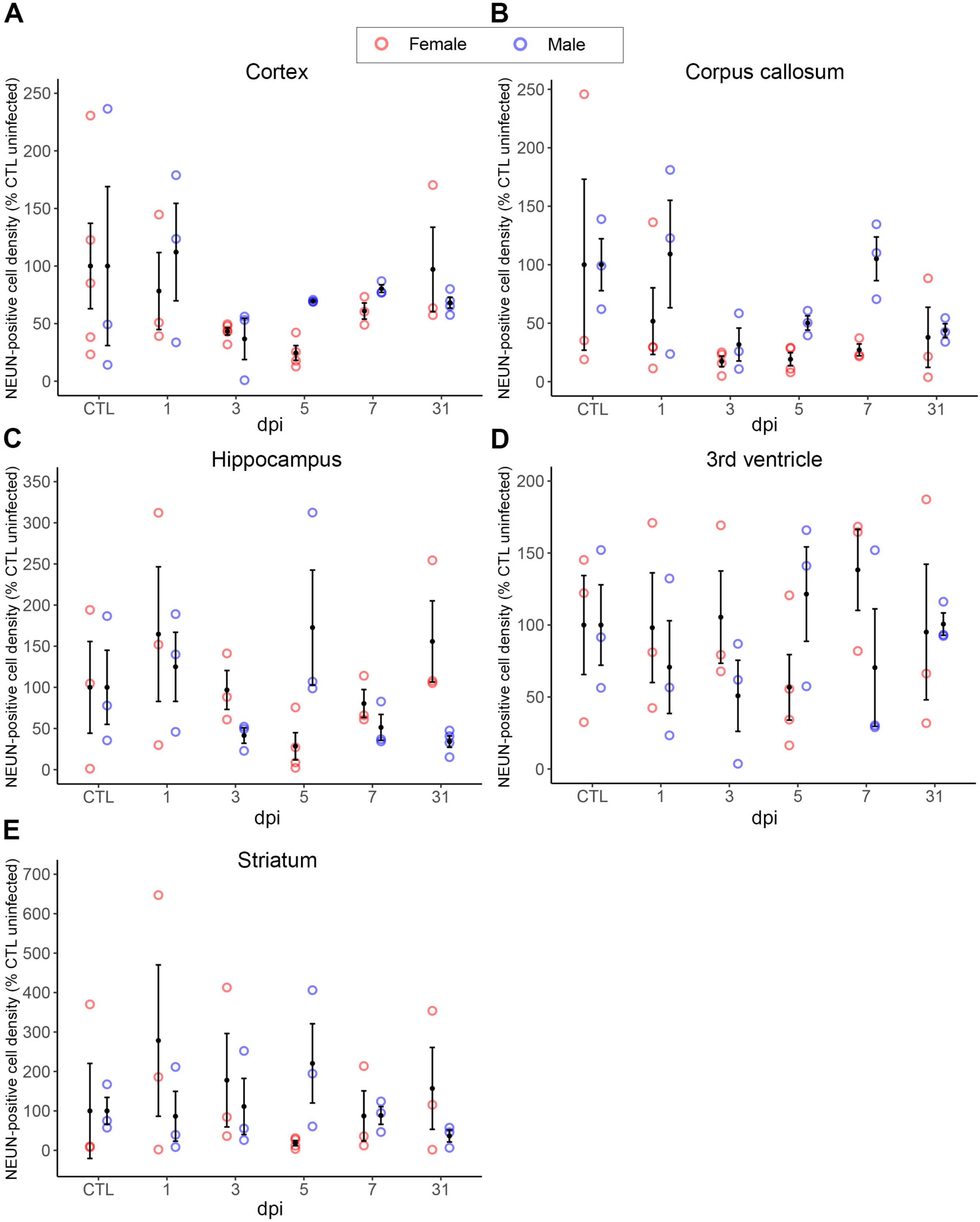
A
B
Female
Male
Cortex
Corpus callosum
NEUN-positive cell density (% CTL uninfected)
dpi
CTL
1
3
5
7
31
C
D
Hippocampus
3rd ventricle
E
Striatum

Figure S5

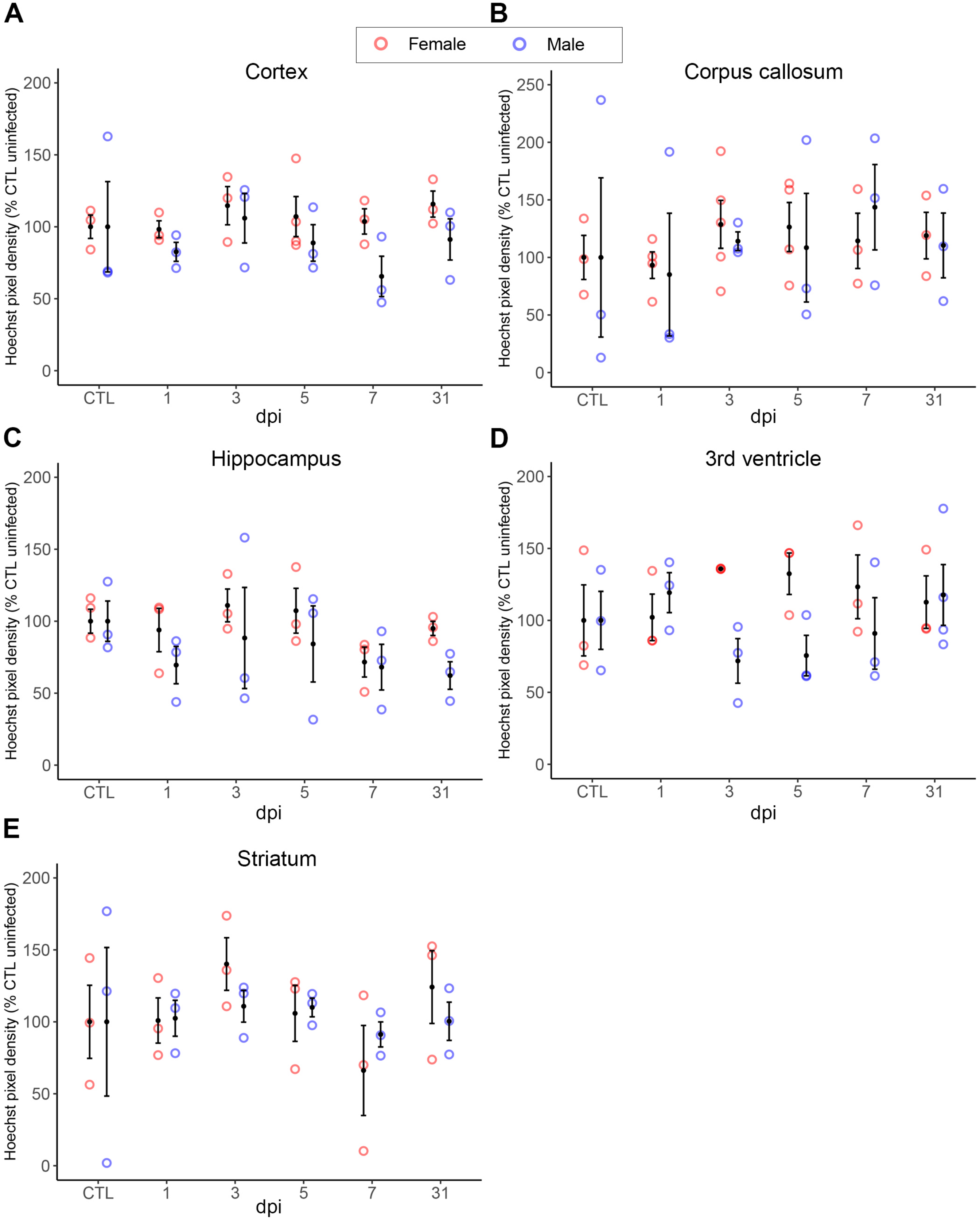
A
B
C
D
E
Female
Male
Cortex
Corpus callosum
Hippocampus
3rd ventricle
Striatum
Hoechst pixel density (% CTL uninfected)
dpi
CTL
1
3
5
7
31
0
50
100
150
200
250

Figure S6

### Supplemental Figure Legends

**Figure S1. Comparison of global thresholding methods.** (A) Three representative confocal micrographs of hamster hippocampi immunolabelled for GFAP. The original image of each is depicted in the top row, with different thresholding methods applied (from top to bottom): Otsu, Kapur's entropy, moment-preserving and minimum error thresholding methods. (B) Peak signal-to-noise ratio (PSNR) from each thresholding method applied to each representative image from (A). Kapur's entropy thresholding elicited the highest PSNR in each image. This figure was adapted from the MSc thesis of MRM ((68); link: https://dspace.library.uvic.ca/items/d7256e07-eee8-4b45-80e7-3c4e4404c299)

**Figure S2. Quantitative analysis pipeline workflow in hamster brain sections labelled with general nuclear marker Hoechst and neuronal marker NEUN.** (Top) Representative tiled and stitched raw confocal immunofluorescence micrographs of Hoechst 33342 (left) and NEUN (right) signals. We first isolate/define the precise region of interest from the full brain slice and determine its area. Next, we improve the signal to noise ratio through background subtraction. Finally, the Hoechst and NEUN signals are converted to a black on white binary format. The pipeline outputs for Hoechst and NEUN (bottom) are normalized to the ROI area, giving signal density for Hoechst, and positive cell density for NEUN as the final read outs. This figure was previously included in the MSc thesis of MRM ((68); link: https://dspace.library.uvic.ca/items/d7256e07-eee8-4b45-80e7-3c4e4404c299)

**Figure S3. GFAP signal density does not significantly change in the hamster corpus callosum, third ventricle, or dorsal striatum following peripheral SARS-CoV-2 infection.** Quantification of GFAP signal intensity from tiled confocal images of full hamster brain sections demonstrate no significant changes in the (A) corpus callosum, (B) third ventricle, or (C) dorsal striatum at any day post-inoculation (dpi) compared to uninfected controls (CTL) in male or female animals. Statistical outcomes are presented in Table S4.

**Figure S4. SOX9-positive cell density is not altered in hamster brains following peripheral SARS-CoV-2 infection.** Quantification of SOX9-positive cells from tiled confocal images of full hamster brain sections demonstrates no significant changes in the (A) cortex, (B) corpus callosum, (C) hippocampus, (D), third ventricle, or (E) dorsal striatum compared to uninfected controls (CTL) in male or female animals. Statistical outcomes are presented in Table S5.

**Figure S5. NEUN-positive cell density is not altered in hamster brains following peripheral SARS-CoV-2 infection.** Quantification of NEUN-positive cells from tiled confocal images of full hamster brain sections demonstrates no significant changes in the (A) cortex, (B) corpus callosum, (C) hippocampus, (D), third ventricle, or (E) dorsal striatum compared to uninfected controls (CTL) in male or female animals. Statistical outcomes are presented in Table S6.

**Figure S6. Hoechst 33342 signal density is not altered in hamster brains following peripheral SARS-CoV-2 infection.** Quantification of Hoechst 33342 (total nuclear marker) signal density from tiled confocal images of full hamster brain sections demonstrate no significant changes in the (A) cortex, (B) corpus callosum, (C) hippocampus, (D), third ventricle, or (E) dorsal striatum compared to uninfected controls (CTL) in male or female animals. Statistical outcomes are presented in Table S7.
